# A novel nuclease is the executing part of a bacterial plasmid defense system

**DOI:** 10.1101/2022.10.06.511153

**Authors:** Manuela Weiß, Giacomo Giacomelli, Mathilde Ben Assaya, Finja Grundt, Ahmed Haouz, Feng Peng, Stéphanie Petrella, Anne Marie Wehenkel, Marc Bramkamp

## Abstract

Cells are continuously facing the risk of taking up foreign DNA that can compromise genomic and cellular integrity. Therefore, bacteria are in a constant arms race with mobile genetic elements such as phages, transposons and plasmids. They have developed several active strategies against invading DNA molecules that can be seen as a bacterial ‘innate immune system’. Anti-phage systems are usually organized in ‘defense islands’ and can consist of restriction-modification (R-M) systems, CRISPR-Cas, and abortive infection (Abi) systems. Despite recent advances in the field, much less is known about plasmid defense systems. We have recently identified the MksBEFG system in *Corynebacterium glutamicum* as a novel plasmid defense system which comprises homologues of the condensin system MukFEB. Here, we investigated the molecular arrangement of the MksBEFG complex. Importantly, we identified MksG as a novel nuclease that degrades plasmid DNA and is, thus, the executing part of the system. The crystal structure of MksG revealed a dimeric assembly through its DUF2220 C-terminal domains. This domain is homologous to the TOPRIM domain of the topoisomerase II family of enzymes and contains the corresponding divalent ion binding site that is essential for DNA cleavage in topoisomerases, explaining the *in vitro* nuclease activity of MksG. We further show that the MksBEF subunits exhibit an ATPase cycle similar to MukBEF *in vitro* and we reason that this reaction cycle, in combination with the nuclease activity provided by MksG, allows for processive degradation of invading plasmids. Super-resolution localization microscopy revealed that the Mks system is spatially regulated via to the polar scaffold protein DivIVA. Introduction of plasmids increases the diffusion rate and alters the localization of MksG, indicating an activation of the system *in vivo*.

## Introduction

Exchange of genetic material is an ubiquitous process. In prokaryotic organisms, horizontal gene transfer (HGT) is the major source of genetic exchange and recombination. Therefore, HGT contributes greatly to the evolution and adaption of prokaryotic organisms (1). HGT is driven by three routes of DNA transfer: virus (phage) infection, conjugation, and transformation. The uptake of foreign DNA is usually associated with a fitness burden and can, in case of phage infection, result in the death of the host cell. Thus, in an attempt to protect their genetic integrity, bacteria are in a permanent arms race with phages and other invading DNA molecules (2). Consequently, bacteria have evolved a multitude of active defense systems, which collectively comprise the bacterial ‘innate immune system’ (3). Among the systems preventing entry of foreign DNA, anti-phage systems are predominant. Classical phage defense strategies include restriction-modification (R-M) systems, CRISPR-Cas, and abortive infection systems (Abi). These systems are usually organized in “defense islands” (4,5). Based on this genomic organization other phage defense systems have been uncovered in recent years. Among those are BREX (6), gasdermins (7), TIR-domain systems (8), dynamin-like proteins (9), and DISARM (10).

So far, less diversity has been observed for plasmid defense systems, with well-known systems such as restriction endonucleases, CRISPR-Cas (11) and prokaryotic argonautes (pAgos) (12) being the main representatives. Recently, a plasmid resistance system centered around a condensin-like ATPase was identified bioinformatically in a large genomic screen (4). The system, termed Wadjet includes four genes (*jetABCD*). JetABC are homologs of condensin complexes MukFEB, respectively. Transplantation of the four genes system into *Bacillus subtilis* led to a reduced transformation efficiency in this heterologous setting (4). In strong agreement with these findings, we have previously shown that in the Gram-positive bacterium *Corynebacterium glutamicum* the MksBEFG system (a Wadjet type I system) is indeed involved in plasmid copy number control (13). Deletion of the condensin subunit MksB (Cg3104) leads to a large increase in the plasmid copy number in *C. glutamicum.* MksB (homolog of JetD) is part of a larger family of ATPase’s that are often involved in chromosome organization and DNA repair (13). The best studied prokaryotic condensin proteins are structural maintenance of chromosome (SMC) and the functionally related MukB protein. While *Escherichia coli* and other gamma proteobacteria encode a MukB dependent system (14), many Gram-positive bacteria as well as eukaryotes use the SMC variant (15). SMC and MukB act in concert with a kleisin protein (MukF in *E. coli* and ScpA in *B. subtilis* and *C. glutamicum)* and a kite protein (MukE and ScpB, respectively) and play a role in chromosome condensation and segregation (16–21). Interestingly, several bacteria harbor homologs of both systems and for long it remained unclear if both work on chromosome organization (22,23). We have shown that while SMC in *C. glutamicum* is involved in chromosome inter-arm cohesion, MksB had no effect on chromosome organization. However, we described an effect of MksB on plasmid copy number, indicating that MksB functions exclusively in plasmid defense and/or control of plasmid copy number (13). These findings are in line with earlier observations in *Mycobacterium smegmatis.* The transformable laboratory strain *M. smegmatis* mc^2^155 carries a spontaneous mutation in an MksB homolog (termed EptC), while presence of EptC leads to restriction of plasmids (24).

Unlike the classical condensin systems the Mks system from *C. glutamicum* contains a fourth subunit, MksG (cg3103). MksG has been predicted to have structural homology with topoisomerase VI (Topo VI). Topo VI is a heterotetrameric complex that identifies DNA crossings and uses ATP hydrolysis to open the DNA and allow strand passing (25)(26). The structures of a topo VI have been solved for the enzyme from *Sulfolobus shibatae* (25) and *Methanocaldococcus jannaschii* (26). Interaction between SMC complexes and topoisomerases is actually common. For example, MukB is required for correct Topo IV localization and activity in Gram negative bacteria (27–30). So far, the mechanism of the condensin-like plasmid defense systems has not been analyzed. We therefore set out to characterize the molecular mechanism of the MksBEFG system from *C. glutamicum* in detail. Our data reveal that MksG is a novel, divalent cation-dependent nuclease that effectively cleaves DNA *in vitro* in an ATP independent manner. Structural analysis of MksG showed that MksG forms a dimer and shares similarities in its catalytic site with the TOPRIM domain of topoisomerase VI. However, MksG differs from TopoVI with respect to the 5Y-CAP and the catalytic tyrosine. MksG activity is not dependent on MksBEF, but ATPase activity of the MksBEF complex is reduced in presence of MksG. The MksBEFG complex is localized to the cell poles in *C. glutamicum.* Single molecule tracking revealed that MksB is mainly confined at the cell pole with an additional slow diffusive population. In contrast, MksG is present in three distinct populations a confined polar population and two dynamic populations across the nucleoid and the cytosol. Presence of plasmid DNA directly changes MksG molecular dynamics. Polar localization of MksG is altered by MksBEF, that in turn colocalize with DivIVA. Depletion of the polar scaffold DivIVA abolishes MksB localization. Thus, a likely scenario is that MksG is held inactive until it encounters the polar MksBEF complex and plasmid DNA. Our data reveal that the plasmid defense mechanism by MksBEFG is executed by the MksG nuclease activity. It also provides evidence for the evolutionary connection for the interactions between MksB/MukB and topoisomerase proteins. Biotechnologically our findings are important for the generation of *C. glutamicum* strains with improved plasmid replication properties.

## Results

### The MksBEFG system is a widespread plasmid defense system

We have shown before that *C. glutamicum* encodes two condensin-like systems. The SMC-ScpA/B system plays a role in chromosome organization and replichore cohesion. The second condensin-like system share homology to the MukBEF system from *E. coli.* In line with the nomenclature proposed by Petrushenko (2011) we termed this second *C. glutamicum* system MksBEFG (22). In addition to the core condensin complex, which comprises an SMC-like ATPase, a kleisin and kite subunits, the Mks system encodes a fourth MksG subunit in its operon. In *C. glutamicum* the four gene operon is located adjacent to the alcohol dehydrogenase (*adhA*) gene on the chromosome (Fig. 1A). The *mks* operon structure (*mksEBEG,* cg3103-cg3106) is conserved also in Gram-negative bacteria harboring this operon, such as *Pseudomonas* species. However, in *P. putida* (KT2440 strain) the *mksB* gene seems to be split while the *P. aeruginosa* (UCBPP-PA14 strain) MksB protein seems to lack the Walker A motif, required for ATP binding (Fig. 1A).

**Figure 1.**
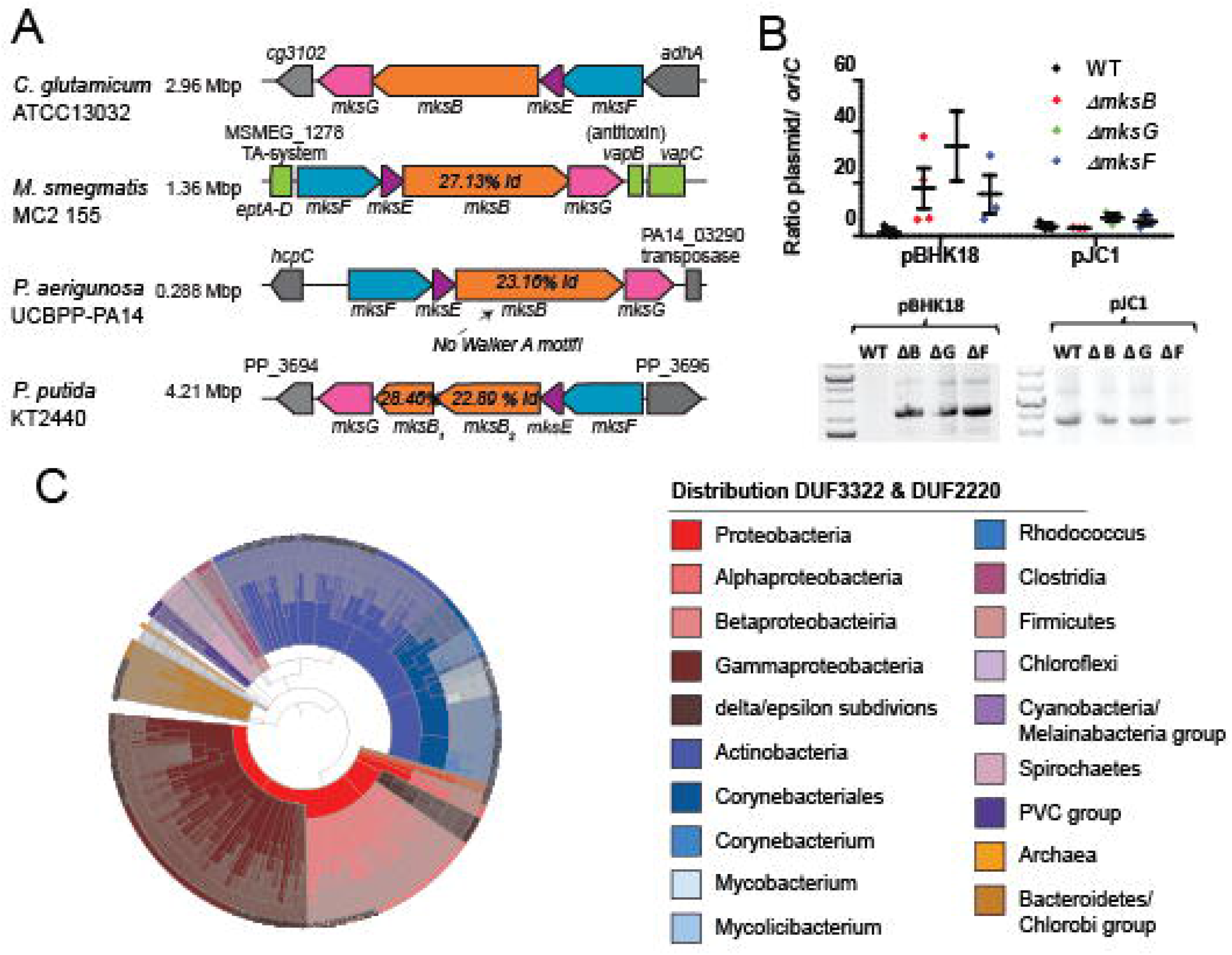
MksBEFG-defense system is widespread. (**A**) Representation of the *mksGBEF-operon* organization from different organisms: (1) *C. glutamicum ATCC13032,* (2) *M. smegmatis mc^2^ 155,* (3) *P. aerigunosa* UCBPP-PA14, (4) *P. putida* KT2440. (**B**) Plasmid copy numbers of low copy (pBHK18) and high copy number plasmids (pJC1) relative to *oriC* numbers per cell, assayed by qPCR. Ratios were compared between *C. glutamicum* WT (MB001), *ΔmksB, ΔmksG, ΔmksF* cells grown in BHI medium with selection antibiotic (mean ± SD, n =3). Plasmids pBHK18 and pJC1 were extracted from *C. glutamicum* WT, *ΔmksB, ΔmksG, ΔmksF* cells grown in BHI medium with antibiotic selection, visualization of extracted DNA on 0.8 % agarose gels. (**C**) Phylogenetic analysis of MksG-like proteins (organized in DUF3322 and DUF2220 domains) using the SMART platform reveals the distribution among Gram-negative and Gram-positive bacteria and archaea.

We have shown before that the MksBEFG system from *C. glutamicum* is involved in plasmid defense and/or copy number control (13). A *mksB* deletion resulted in a significant increase of low-copy plasmids (pBHK18), but not medium-copy plasmids (pJC1) (13). Here, we show a detailed analysis of the MksBEFG complex and of its role in plasmid defense. Similar to the deletion of *mksB,* null alleles of *mksG* and *mksF* have a significant effect on the copy number of the pBHK18 plasmid, but not of pJC1 (Fig. 1B). Isolation of plasmid DNA from the different strain backgrounds revealed that deletion of either *mksB, mksF* or *mksG* leads to an increase in plasmid yield, in line with the qPCR data. Thus, the presence of all three major subunits of the Mks system, including the so far uncharacterized subunit MksG, is required for an effective plasmid defense.

MksG comprises two domains of unknown function (DUF3322 and DUF2220). A domain organization analysis using the SMART platform (31) revealed that there are 1,366 proteins with the same domain architecture (DUF3322 plus DUF2220) in bacteria and archaea (Fig. 1C). MksG-like proteins are encoded not only in Corynebacteriales, but in a wide variety of bacterial clades (47.8 % proteobacteria, 37.8 % actinobacteria, 4.2 % bacteriodetes, 2.5 % spirochaetes, 2.2 % firmicutes and 5.5 % others). The Mks system was also identified in some archaea. Interestingly, MksG-like proteins are not found in all members of a family and hence it is likely that the gene was spread by HGT, as suggested earlier (4).

### MksG reveals a topoisomerase fold and exhibits ion-dependent nuclease activity

To get insights into the function of MksG we determined its structure by combining low resolution X-ray data at 4.6 Å resolution with high confidence Alphafold2 (AF2) models (TableS1). MksG is formed by two domains, an N-terminal (N-ter) elongated domain formed of mostly α-helices as well as an elongated 3 stranded ß-sheet, and a C-terminal (C-ter) domain with a central ß-sheet, flanked by α-helices (Fig. 2A). The C-ter domain forms a dimer in the crystal by extending the central ß-sheet. The N- and C-terminal domains are connected by a flexible hinge region (Fig. 2AB). A structural homology search using DALI on the AF2 database revealed that the C-ter domain is homologous to the topoisomerase VI (Topo VIA) domain of the Rec12/Spo-11 meiotic recombination factor. Topo VI belongs to the topoisomerase II family members that function as hetero-tetramers of two A-subunits, containing the TOPRIM and 5Y-CAP catalytic domains responsible for DNA cleavage as well as two ATPase B-subunits that are responsible for DNA positioning and movement (26,32).

**Figure 2.**
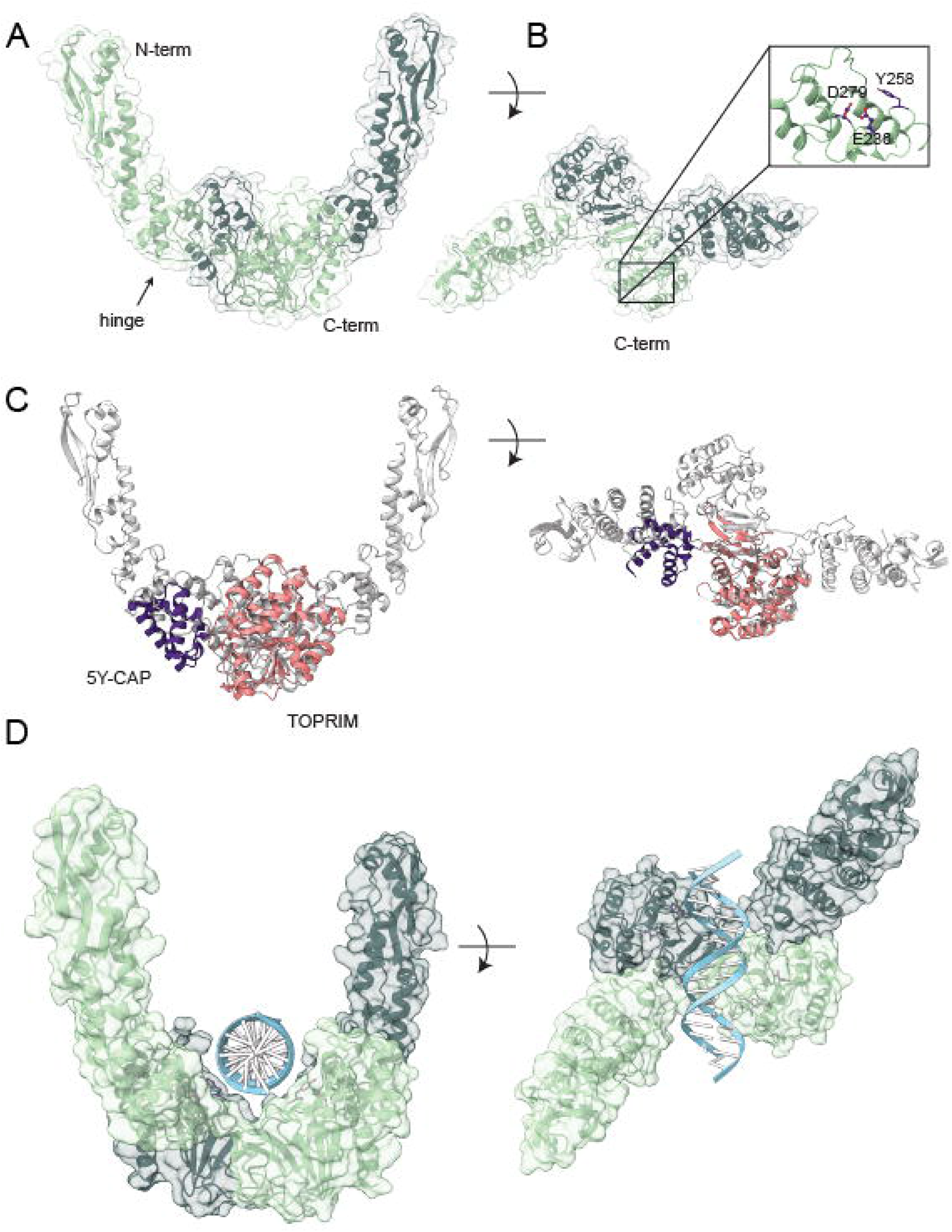
Structural characterization of MksG. (**A**) Cartoon and surface representation of the dimer of MksG. Each monomer is shown in light and dark green respectively. The flexible hinge region for one monomer is indicated by an arrow. (**B**) The C-terminal domain is shown as a cartoon and the position of the predicted active site residues is highlighted. The insert shows the Mg^2+^ binding residues E236 and D279, as well as the putative active site Y258. (**C**) Superposition of Topo VIA from *M. jannaschii* (PDB code 1d3y). The TOPRIM domain (in pink) superposes well with the C-terminal domain of MksG with an RMSD between 52 pruned atom pairs is 1.074 angstroms (calculated using matchmaker of ChimeraX). The 5Y-CAP domain of Topo VIA is shown in purple, and overlapping with the lower part of the N-terminal domain of MksG. (**D**) A model of a double stranded DNA molecule was positioned above the two active sites of MksG to visualize a possible DNA binding mode.

Based on the obtained structural data we expected MksG to have nuclease activity. Based on our observation that loss of MksG leads to plasmid enrichment in *C. glutamicum* (Fig. 1 and (13)) we used plasmid DNA for nuclease experiments. Plasmid DNA is present in three conformations: open-circular (OC) (nicked), linear (L) and super-coiled/ closed-circular-conformation (CCC). The untreated plasmid was found to be predominantly in the closed circular conformation. Addition of MksG led to an increase in the nicked (OC) and linear conformation, while the supercoiled fraction decreased (Fig. 3). Longer incubation (more than 1 h) lead to a degradation of the plasmid DNA (Supplementary Figure S2), indicating that MksG is indeed an active nuclease.

**Figure 3.**
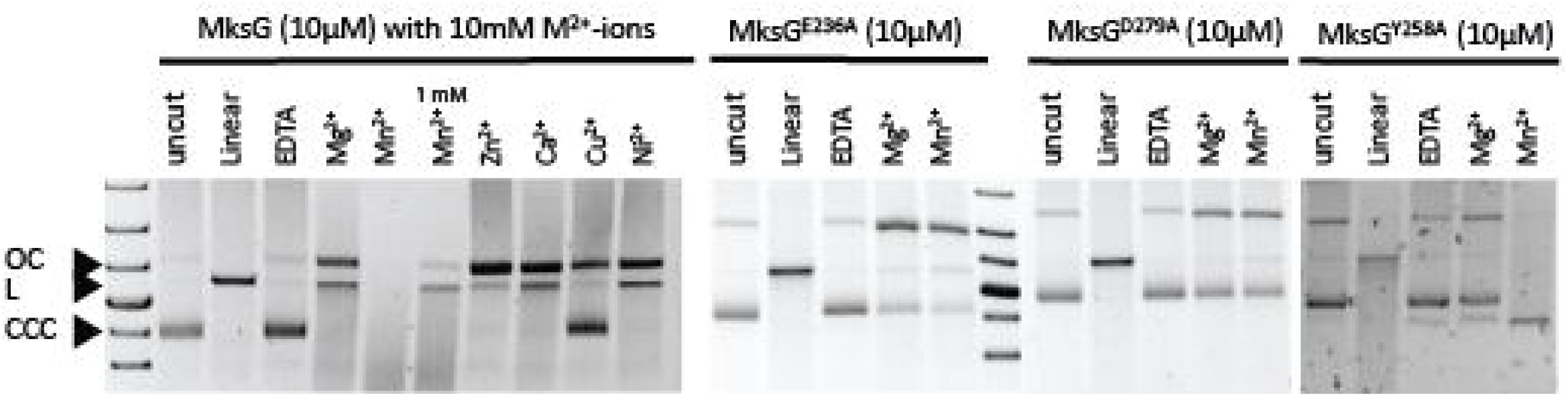
MksG is the active nuclease subunit of the defense system. Nuclease activity assay of MksG, 10μM protein were incubated 1h at 30°C with 250 ng plasmid DNA (pBHK18, 3,337 bp) and 10 mM different divalent Metal-ions (second lane with Mn^2+^ only 1 mM). Reaction was stopped by adding 6x purple loading dye (NEB) and boiling samples 5 min at 90°C. DNA was separated on an agarose gel in TAE buffer. Assays for mutants (MksG^E236A^, MksG^D279A^, MksG^Y258A^) were incubated 3h at 30°C.

DNA cleavage occurs via a general acid-base mechanism where a general base deprotonates the active site tyrosine hydroxyl, allowing the oxyanion to attack the scissile phosphate of the DNA and a divalent metal, generally Mg^2+^ is required to stabilize the scissile phosphate (33). Sequence alignments and the structural superposition with the Topo VI A subunit of *Methanocaldococcus jannaschii* (26) show a conserved position for the magnesium binding residues essential for activity (Fig. 2B). Therefore, we investigated ion dependency of MksG activity and thus tested the effect of different divalent metal ions on the DNA cleaving activity of MksG (Fig. 3). The highest nuclease activity was observed for 10 mM Mn^2+^-ions. Addition of the metal ion chelator EDTA completely abolished MksG nuclease activity. Other divalent cations such as Ni^2+^, Zn^2+^ and Ca^2+^ also catalyze a weak, but detectable nicking activity, while Cu^2+^ ions do not support activity (Fig. 3).

To corroborate the ion binding-site we mutated two of the amino acid residues, E236 and D279, predicted by sequence alignment (Supplementary Figure S3) and structural analysis to be essential for the ion binding, to alanine. We expected that the mutants would not be able to bind the ion ligand (Mg^2+^/Mn^2+^) and therefore lose the ability to cleave DNA. In line with our hypothesis the two MksG mutants (MksG^E236A^, MksG^D279A^) had drastically impaired nuclease activity (Fig. 3).

Structural comparison of MksG with TopoVIA shows that the TOPRIM domain dimerizes, in the same way in both proteins (Fig. 2), to form a DNA binding groove and allow for double stranded DNA breaks. Interestingly, in MksG only the TOPRIM domain is conserved. It is thus not clear where the catalytic tyrosine, normally found in the 5Y-CAP domain, comes from. The N-terminal domain of MksG is positioned in a similar manner to the 5Y-CAP domain in Topo VIA with respect to the TOPRIM domain (Fig. 2C). However, no tyrosine residues are present in the vicinity that could have a catalytic role. The only tyrosine within reasonable distance for a putative catalytic role is Tyr258 (Fig. 2C), located within the TOPRIM domain. Interestingly from the DALI search on the AF2 database, several eukaryotic homologs of the Spo11 family have a conserved Tyr in the same position. The tyrosine engages in a covalent bond formation with the 5’phosphate of the DNA. This covalent enzyme-DNA complex can be religated in topoisomerase activities (34–36). To test whether Y258 is involved in the DNA cleaving activity of MksG, we replaced Y258 by an alanine (Y258A mutant). Nuclease activity assays revealed that indeed the Y258A mutant has a drastically reduced cleavage activity (Fig. 3). However, a residual activity was still observed.

A modelled DNA molecule could be placed into the putative binding cleft formed by the MksG dimer and the distance between the two active sites could be compatible with double stranded DNA breaks (Fig. 2D). The elongated N-terminal domain may also serve to stabilize DNA-binding. This part resembles the transducer domain of the B subunit of topoisomerases that links the Bergerat ATPase domain with the 5Y-CAP of the A subunit. Importantly, MksG lacks any ATP binding domain. The structural analysis together with the catalytic activities provide strong evidence that MksG is indeed a bona fide Topo VIA homolog.

### MksB ATPase is stimulated by complex formation and DNA binding

MksG is encoded in an operon with the condensin-like MksBEF complex. Since deletion of MksG has the same effect on plasmid stability as the MksB or MksF deletions (Fig. 1), these proteins likely act as a complex in vivo. The *mksB* gene encodes for an 1,111 amino acid condensin-like polypeptide with a calculated molecular weight of 126.58 kDa. The sequence analysis reveals that MksB shares the five-domains architecture with other condensins such as SMC and MukB ((1) the conserved amino-terminal (N-terminal) Walker-A motif; (2) coiled-coil region; (3) hinge-domain; (4) coiled-coil domain; (5) conserved carboxy-terminal (C-terminal) Walker-B and signature-motif) (16,22,37,38) suggesting that the MksB from *C. glutamicum* also belongs to the ABC class of ATPases (Supplementary Figure S4). Even though the structure of condensins is highly conserved the sequence identity is rather low (~25 %) (Fig. 1, Supplementary Figure S4).

In order to understand and characterize the molecular complex, the four proteins were heterologously expressed with a poly-His-Tag in *E. coli* and purified in via several chromatography steps (see material and methods). All proteins were purified to homogeneity and used for the *in vitro* assays (Fig. 4A). Although MksF and MksE could be purified individually, co-purification of both proteins stabilized the proteins against degradation during the isolation process.

**Figure 4.**
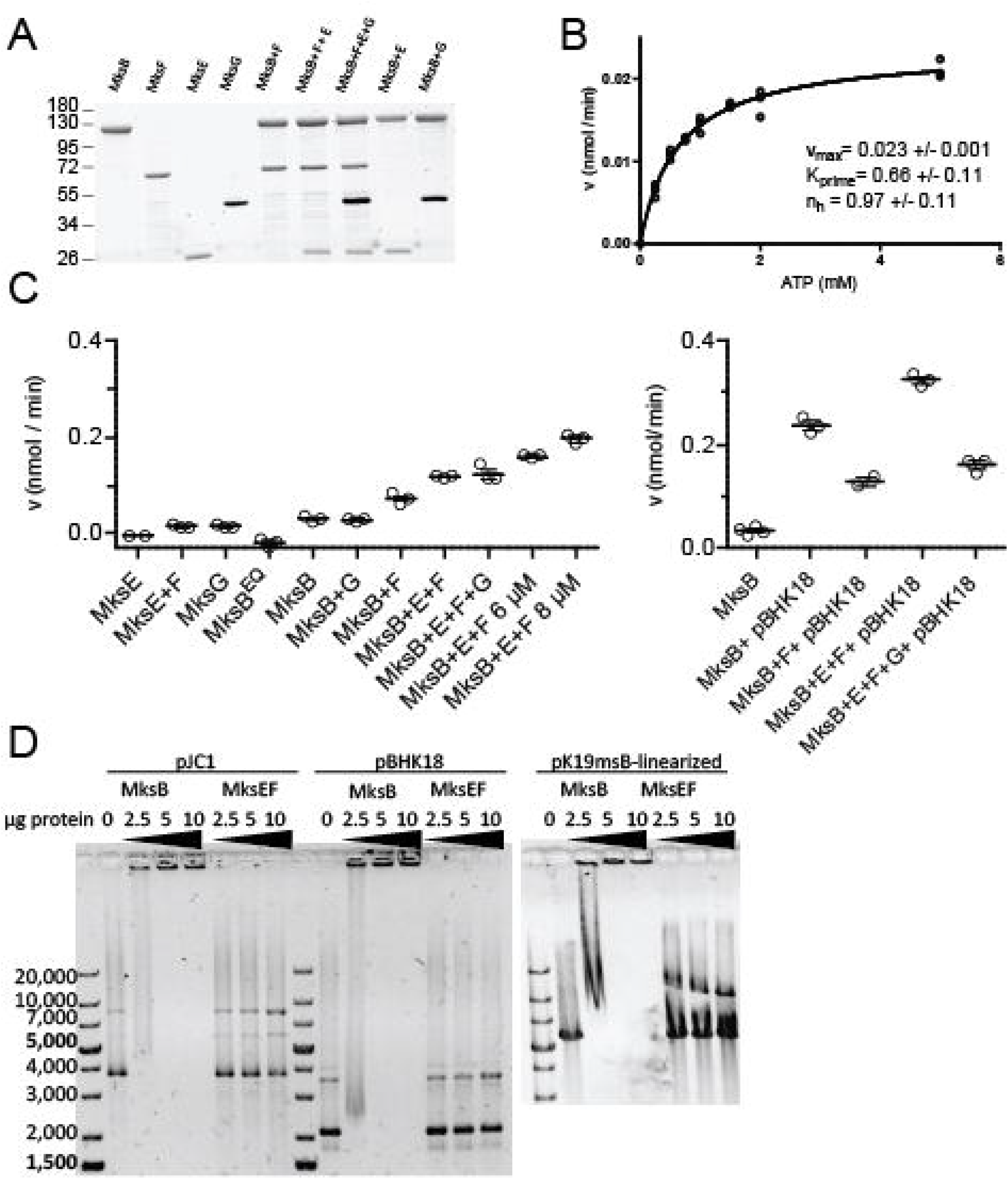
MksB ATPase activity and DNA binding *in vitro.* (**A**) SDS-PAGE (stain-free) of purified Mks-proteins after ATPase assay, indicating highly purified Mks proteins. (**B**) MksB (4 μM) ATPase activity following Michaelis Menten kinetics at different ATP concentrations (0-5 mM). (**C**) MksB (4 μM) ATPase activity analysis in combination with the other subunits, MksB^EQ^-mutant and plasmid pBHK18. 4 μM of each subunit were applied, unless stated otherwise, plasmid DNA was 50ng/μl. ATPase measurements were taken in a time-course of 3h, no substrate controls and ATP autohydrolysis were subtracted from values. Each data point represents one measurement, the mean is shown as line with standard error, n=3). (**D**) MksB binds circular and linear plasmid DNA independent of Mg-ATP. MksEF are not able to bind to DNA on their own. Electrophoretic mobility shift assays of MksB, MksEF were preincubated with 200 ng circular (pJC1, 6,108 bp; pBHk18, 3,337 bp) and linear (pK19mobsacB, 5,722 bp) plasmid DNA.

MksB revealed a low basal ATPase activity that followed Michaelis-Menten kinetics (Fig. 4B). For this, 4 μM of MksB were incubated with different concentrations of Mg^2+^-ATP (0-5 mM) (Fig. 4B). A Hill-coefficient of nh=0.97 was determined, confirming that MksB ATPase activity is not cooperative. The calculated vmax value was 0.023 nmol/min and is quite low compared with other MukB-like ATPases (39). As expected, in the Walker B hydrolysis deficient mutant MksB^E1042Q^ ATPase activity was impaired (Fig. 4C). However, we observed in a titration assay that the activity values calculated negative values (subtracting no substrate and ATP autohydrolysis values) suggesting that the mutant MksB^EQ^ is able to bind ATP that is the protected from auto-hydrolysis resulting in negative values. In line with data described for the MukB/MukF interaction addition of equimolar concentrations of the kleisin subunit MksF and the kite subunit MksE stimulated the ATPase activity of MksB (Fig. 4C). Further rise of MksEF concentration (6 μM and 8 μM each) lead to further increase of MksB’s activity. Addition of equimolar amount of MksG to MksBEF had no impact on ATPase activity. Furthermore, the addition of plasmid DNA (pBHK18) led to an increase of MksB’s activity (Fig. 4C). The maximum ATPase activity that we measured was with equimolar amounts of MksBEF and plasmid DNA (0.32 nmol/min). Interestingly, the addition of MksG to MksBEF and plasmid DNA led to a significant decrease of activity (0.16 nmol/min), likely because of the nuclease activity of MksG that might reduce the DNA concentration within the assay. The MksBEFG system from *C. glutamicum* displays significant differences with the MukBEF complex, where addition of the kite subunit decreases ATPase activity, suggesting this activity could be differently regulated in the two systems.

MksB alone is able to bind and shift DNA in gel electrophoretic mobility shift assays (EMSA) (Fig. 4D). Binding of MksB to linear, supercoiled or nicked-plasmid DNA was ATP-independent. MksEF alone are not able to bind and shift DNA. These data suggest that MksB is the DNA-binding subunit in the MksBEFG complex, similar to the well characterized MukBEF complex. *In vitro* we were not able to identify any selectivity of MksB binding to DNA. However, we speculate that *in vivo* the MksBEFG complex likely identifies incoming plasmid DNA. The precise mechanism why and how the *C. glutamicum* MksBEFG system reacts to low-copy number plasmids and not to others remains to be analyzed.

### MksF links MksG to the MksBEF complex

The complex assembly for the *E. coli* MukBEF is relatively well known (40,41). However, it remained unclear how the MksBEFG complex is assembled. In particular the localization of the nuclease subunit MksG was unclear. Therefore, we used Bio-Layer Interferometry (BLI) to analyze how the subunits interact with each other. For this we biotinylated each protein to test different conformations and immobilized always one subunit at a time with Streptavidin (SA) biosensors. Having one bait protein immobilized to the sensor, association with the analyte protein (different concentrations were tested) was performed. Several combinations did not show interactions. We tested immobilization of MksF with MksB and MksE as analytes, MksG immobilized with MksB and MksE as analytes and immobilized MksB and MksE as analyte. In all these combinations no protein-protein interaction was observed via BLI (data not shown). Successful BLI interactions where shown for the kleisin and kite subunits MksF and MksE (Fig. 5A). This interaction was expected since the kite and kleisin subunits in other MukBEF-like systems also interact. We immobilized MksE (25μg/ml) and titrated with MksF (1-5 μM) (Fig. 5). As expected from previous chromatography experiments with MksEF copurification, we observed the strongest interaction between MksE and MksF with a K_D_ (equilibrium dissociation constant) value of 2.17 μM (Table 1). At the highest concentration of MksF (5 μM), 1:1-binding of MksE:MksF could not be observed anymore. This suggests that there is more than one binding site of the analyte to the bait and heterologous binding occurs. This implies that one MksF molecule binds to two immobilized MksE molecules, as it has been described for MukF-MukE or ScpA/ScpB interaction.

**Figure 5.**
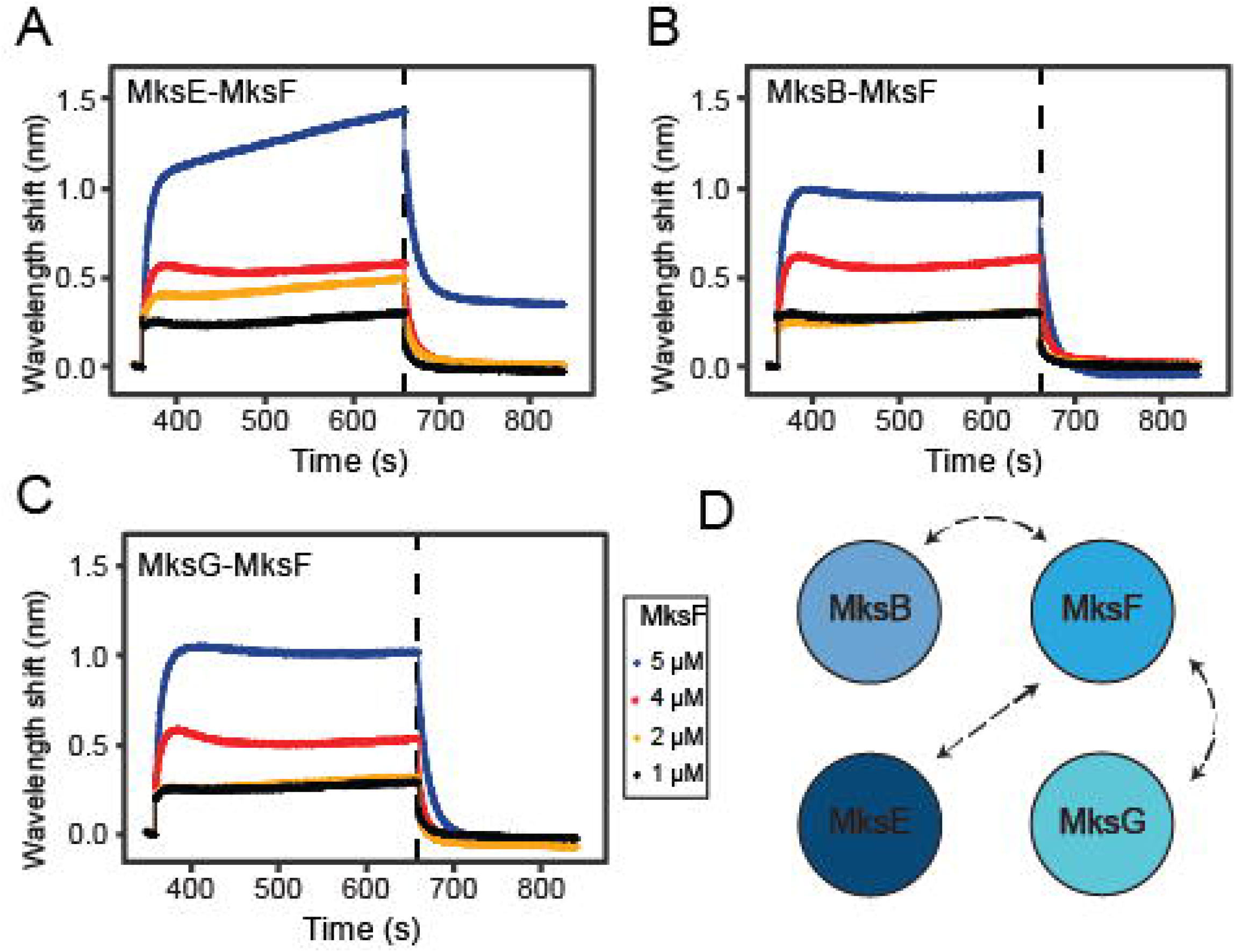
MksF is the interaction hub of the MksBEFG complex. Bio-Layer interferometry of (**A**) immobilized biotinylated MksE (25 μg/ml) with MksF (1-5 μM) (**B**) immobilized biotinylated MksB (50 μg/ml) with MksF (1-5 μM) and (**C**) immobilized biotinylated MksG (100 μg/ml) with MksF (1-5 μM). Initial Baseline was generated for 30s, loading of the bait was performed for 300s, a second baseline was generated for further 30s, association with the analyte was performed for 300s and final dissociation was performed for 180s. (**D**) Interaction map of the Mks subunits based on BLI analysis.

**Table 1.** Protein-protein interaction among complex.

| Interaction partners | $K_D$ [$\mu M$] | $k_a$ [$M^{-1} s^{-1}$] | $k_{a_{error}}$ | $k_d$ [$s^{-1}$] | $k_{d_{error}}$ | $\chi^2$ | $R^2$ |
| --- | --- | --- | --- | --- | --- | --- | --- |
| MksE-MksF | 2.17 | $7.42 \cdot 10^4$ | $0.48 \cdot 10^4$ | 0.16 | 0.0027 | 1.9 | 0.98 |
| MksB-MksF | 4.94 | $2.78 \cdot 10^4$ | $0.16 \cdot 10^4$ | 0.138 | 0.0018 | 2.28 | 0.99 |
| MksG-MksF | 6 | $2.25 \cdot 10^4$ | $0.16 \cdot 10^4$ | 0.136 | 0.002 | 3.041 | 0.99 |

**Table 2.** BLI Advanced Kinetics Protocol.

| Step type | Sample type | Position | Duration (s) |
| --- | --- | --- | --- |
| Initial Baseline | Buffer | Tube | 30 |
| Loading of the bait | Biotinylated Protein | Drop | 300 |
| Baseline | Buffer | Tube | 30 |
| Association | Analyte (MksF) | Drop | 300 |
| Dissociation | Buffer | Tube | 180 |

Furthermore, we immobilized MksB (50μg/ml) and measured interaction with MksF (1-5 μM) (Fig. 5B). Here, we observed interaction with a K_D_ value of 4.94 μM (Table 1) and no apparent heterologous binding. Testing further possibilities, we found that immobilizing MksG (100 μg/ml) and using MksF as analyte we observed clear interaction. We determined the K_D_ value of 6 μM as the highest value, resulting in the lowest affinity of the complex. Most importantly, we could show the interaction of MksG with the kleisin subunit MksF (Fig. 5C). This was somewhat surprising, since in *E. coli* the ATPase MukB interacts physically with the topoisomerase IV. However, in the MksBEFG system the topo VI-like MksG rather interacts with the kleisin subunit. All kinetic parameters from the biolayer inferometry experiments are summarized in Table 1.

From these data, we conclude that MksF serves as an interaction hub between the central complex (MksBEF) and the novel nuclease subunit MksG.

### Localization of the MksBEFG complex *in vivo* and interaction *in vitro*

Previously, we reported that MksB co-localizes with the polar scaffold protein DivIVA (13). Next, we wanted to analyze and compare the localization of each subunit of the complex. Therefore, each subunit was genetically fused with the sequence of Halo-Tag and inserted into the native locus in the genome, to preserve physiological protein levels. Importantly, in-gel fluorescence revealed that all fusions were expressed as full-length proteins. We observed no major degradation of the fusions (Supplementary Figure S5), suggesting that the constructs were well suited for microscopic analyses. We had shown before that MksB-mCherry localized to the cell poles and may interact with the polar scaffold protein DivIVA (13). MksB-Halo localized as expected to the cell poles and septa in *C. glutamicum* (Fig. 6A). To test if this localization is DivIVA or cell pole dependent, we constructed a CRISPRi depletion strain in which we could downregulate the expression of the essential DivIVA protein (42). After DivIVA depletion cells lose their rod-shaped morphology and the MksB localization gets random (Supplementary Figure S6).In wild type cells, MksE-Halo and MksF-Halo exhibited the same polar localization was observed as for MksB-Halo (Fig. 6BC), as expected based on the strong interaction of these proteins within the MksBEF complex. Despite MksBEF localizing similarly to the cell poles, we found that MksG-Halo localization to the cell poles was less pronounced (Fig. 6D). Rather, a large proportion of MksG localized as a cytosolic molecule that does not form strong polar foci, but a septal enrichment remained obvious (Fig. 6D). Localization of MksG remained less polar in presence of a plasmid (pBHK18) (Fig. 6E), however deletion of *mksB* had a significant effect on MksG localization. In the *mksB* null allele background MksG localized more polar and less cytoplasmic (Fig. 6F). This effect became even more pronounced when a plasmid was present in the *mksB* deletion background (Fig. 6G). Here, MksG became strongly polar localized. Fluorescent micrographs showed cell with large assemblies of MksG-Halo close to the cell poles.

**Figure 6.**
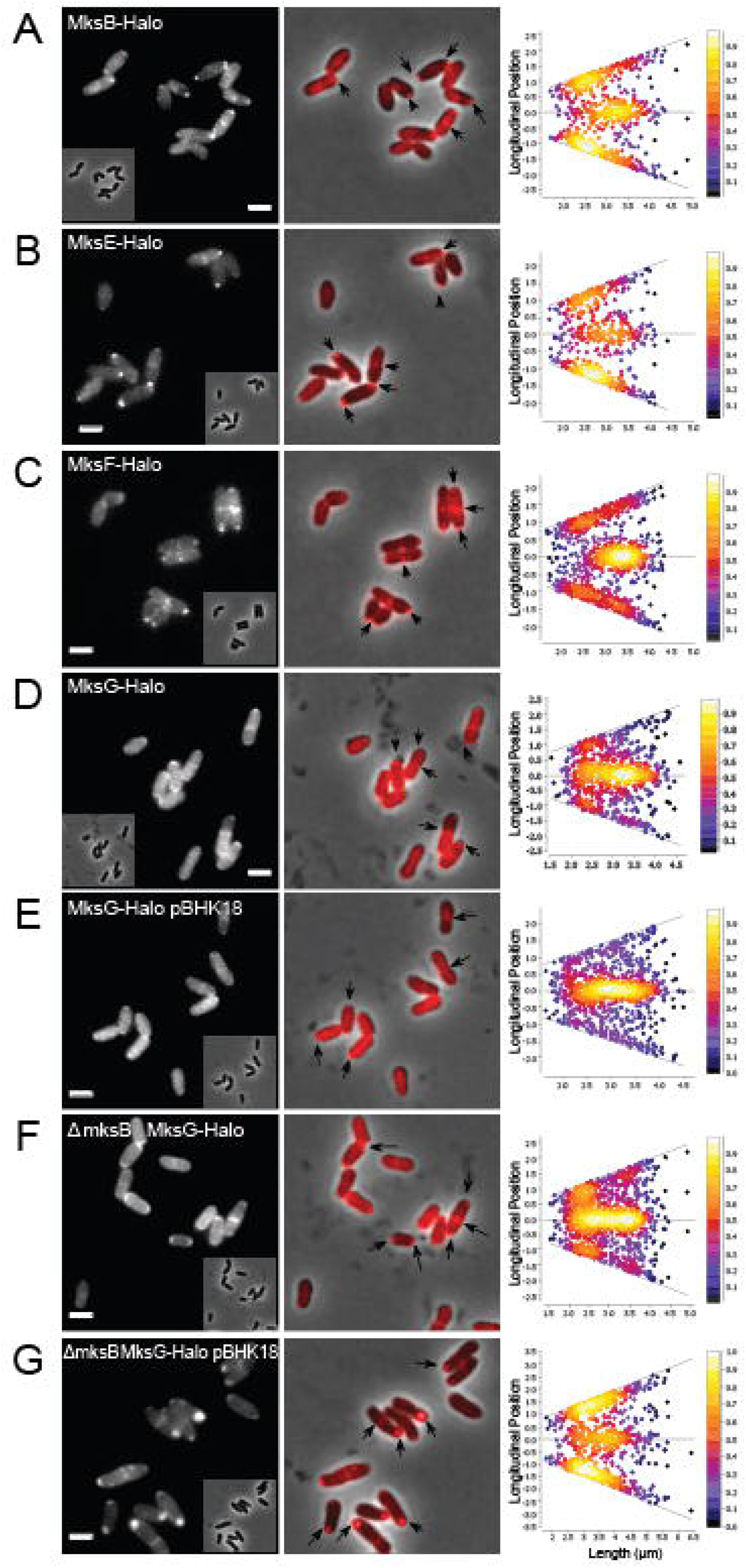
MksBEFG localization *in vivo*. Epifluorescence microscopy images of fusion proteins with Halo-Tag labelled with TMR as a ligand (**A**) of CMG015 cells (number of cells, n=549) (MksB-Halo) (**B**) of CMG013 cells (n=536) (MksE-Halo) (**C**) of CMG014 cells (n=863) (MksF-Halo) (**D**) of CMG012 cells (n=633) (MksG-Halo) (**E**) of CMG032 cells (n=574) harboring pBHK18 plasmid (MksG-Halo + pBHK18) (**F**) of CMG018 cells (n=603) (Δ*mksB* MksG-Halo) (**G**) of CMG034 cells (n=510) harboring pBHK18 plasmid (*ΔmksB* MksG-Halo + pBHK18), scale bar 2 μm. Demographs show the fluorescence maxima distribution along the cell axes sorted by the length of the cell.

These data indicate that the MksBEFG system is spatially confined to the cell poles in *C. glutamicum.* However, MksG seemed to be more mobile and less well constricted to the cell poles. This changed dramatically, when MksB was absent, suggesting that the more diffusive behavior of MksG depends on the presence of MksB.

### MksG dynamics depend on MksB and plasmid DNA

To further characterize and understand the relationship between MksG and MksB localization, we used single molecule localization microscopy (SMLM) in combination with single particle tracking (SPT). SPT experiments were performed at 20 ms exposure time. We aimed to understand how the dynamics of the MksG nuclease change in the presence or absence of plasmid DNA. For this analysis, we used MksG-Halo and MksB-Halo tag fusions stained with TMR dye as the fluorescent ligand as described above.

First, we compared the dynamics of MksB-Halo and MksG-Halo separately (Fig.7AB). The first striking observation was the apparently much faster dynamics of MksG-Halo compared to MksB-Halo. Using the mean-squared displacement (MSD) as a quantitative indicator of the averaged diffusion speed we showed that the diffusion coefficient of MksG-Halo (D= 0.241 μm^2^ s^-1^) was 4.4 x larger than for MksB-Halo (D = 0.055 μm^2^ s^-1^) (Fig. 7B). The slope of the linear fit served as a good estimation for the diffusion coefficient for simple Brownian motion of single population that does not undergo changes in motion. Importantly, the dynamics of MksG was altered in the absence of MksB. In a strain background were *mksB* was deleted the dynamics of MksG decreased (D = 0.164 μm^2^ s^-1^) (Fig. 7AB). However, addition of plasmid DNA accelerated MksG dynamics (D = 0.264 μm^2^ s^-1^). MksG dynamics was also increased when plasmid DNA was present in absence of MksB (D = 0.219 μm^2^ s^-1^), indicating that MksG may be able to bind to plasmid DNA *in vivo* without the interaction of MksB. Presence of plasmid DNA in cells lacking MksB still increased the dynamics of MksG (D = 0.219 μm^2^ s^-1^), albeit to a lesser extent.

**Figure 7.**
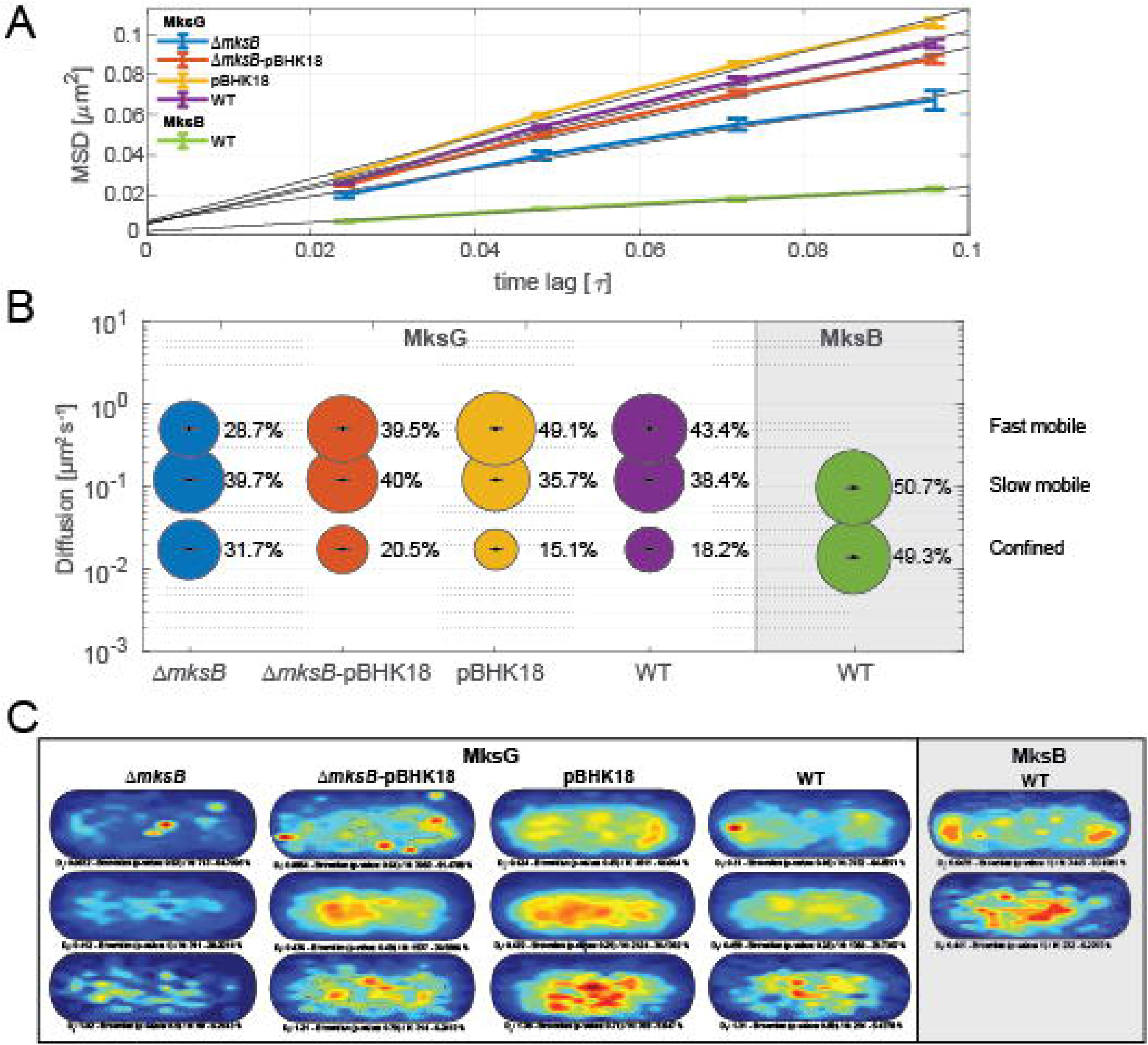
MksG dynamics depend on MksB and plasmid DNA. (**A**) Plot of the mean-squared displacement of the indicated MksG-Halo (in different background strains) / MksB-Halo fusions over time. (**B**) Bubble plot showing single-molecule diffusion rates of the indicated MksG-Halo (in different background strains)/ MksB-Halo fusions. Populations were determined by fitting the probability distributions of the frame-to-frame displacement (jump distance) data of all respective tracks to a three components model (fast mobile, slow mobile, and confined protein populations). (**C**) Confinement heat maps of specified MksG-Halo (in different background strains) / MksB-Halo fusions, indicating likeliness of presence of confined molecules from low (blue) to high (red) in an averaged cell. Strains used for SPT analysis were MksB-Halo (CMG015), MksG-Halo (CMG012), MksG-Halo + pBHK18 (CMG032), *ΔmksB* MksG-Halo (CMG018), *ΔmksB* MksG-Halo + pBHK18 (CMG034),

Given that proteins dynamics do not usually originate from a single population that moves via Brownian motion, but rather comprise separate subpopulations, each defined by its own parameters, we further proceeded to perform a jump distance (JD) analysis. Thus, we were able to identify three distinct populations (fast mobile, slow mobile and confined) for MksG-Halo (WT) and two (slow mobile and confined) for MksB-Halo (WT) (Fig. 7B). Strikingly, MksG has a large fast-mobile population (D = 0.509 μm^2^ s^-1^ / 43.4 %), which is instead lacking in MksB dynamics. This fast-mobile population is decreased in a strain with *mksB* deletion (28.7 %) and increased in the presence of plasmid DNA (49.1 %). Both MksB and MksG exhibit slow mobile and confined protein subpopulations which are characterized by similar, independently fitted, diffusion rates (Slow mobile: DMksG = 0.123 μm^2^ s^-1^, DMksB = 0.0982 μm^2^ s^-1^/ Confined: DMksG = 0.0176 μm^2^ s^-1^, DMksB = 0.0141 μm^2^ s^-1^) (Fig. 7B). In line with the localization studies (Fig. 6) both MksB and MksG confined subpopulations are enriched at the cell poles, while all the remaining subpopulations are characterized by a rather homogeneous localization (Fig. 7C).

We could further observe changes in MksG-Halo dynamics in cells lacking MksB and/or harboring the pBHK18 plasmid. In detail, cells lacking MksB (*ΔmksB)* were characterized by a ~13 % shift from the fast moving to the confined subpopulation (Fig. 7B). On the opposite, cells that harbor the low-copy plasmid pBHK18 were characterized by an increase in the fast-moving subpopulation at the cost of a generalized decrease in both the confined and slow-moving subpopulations (Fig. 7B). To complete the picture, we analyzed the dynamics of MksG-Halo in absence of MksB but presence of pBHK18. Here, we observed a quasi-wild type phenotype, indicating that the previously noted increase in MksG dynamicity due to addition of plasmid is maintained in the *mksB* deletion mutant (Fig. 7B).

Finally, we analyzed the spatial distribution of the three MksG protein subpopulations in the above-mentioned genetic backgrounds. The most confined MksG populations were localized to the cell poles (Fig. 7C). When plasmid DNA was present, the confined MksG population was also found in larger patches around the site of septation. Importantly, absence of MksB greatly diminishes the polar localization of the confined MksG populations, thereby indicating that the MksB affects MksG polar localization (which in turn was shown to depend on DivIVA) (43). The dynamic populations of MksG are dispersed within the cytoplasm (or across the nucleoid) in *C. glutamicum* (Fig. 7C). This proportion is largely increased in the presence of plasmid.

In summary, SMLM data suggest that the MksBEFG system consist out of a confined, polar population and a more diffusive population. MksG dynamics is modulated by MksB and, importantly, by the presence of plasmid DNA.

## Discussion

Bacteria have evolved sophisticated complexes to protect their genomic integrity from incoming foreign DNA. The protection systems include those that act against plasmid DNA. In recent years several new defense machineries in addition to the restriction endonucleases, CRISPR, and argonaute have been described. One of these is the Wadjet system that is composed of a condensin-like complex (4). We have shown earlier that *C. glutamicum* harbors a Wadjet type I system that we termed MksBEFG (43). The Sorek group has transplanted a Wadjet operon into *B. subtilis* and could show that the recombinant strain has drastically reduced transformation frequencies (4). They observed the strongest effect for type I wadjet system. We have shown in *C. glutamicum* that deletion of the condensin subunit MksB leads to a massive enrichment of plasmid copy number. In particular intriguing was the observation of a polar localization of MksB-mCherry in *C. glutamicum* (13). Despite these observations, the data on the molecular function of the Wadjet systems remained sparse. We have therefore set out to analyze the biochemical and cell biological features of the corynebacterial MksBEFG complex in more detail.

All condensin like complexes exhibit a regulated ATPase activity that is linked to the function of these complexes on DNA organization (44). A unifying activity of all SMC complexes is the formation of DNA loops (loop-extrusion) (44–47). This activity on the DNA substrate requires ATPase activity and without a full ATPase cycle SMC complexes do not entrap topologically their DNA substrates. In contrast to many other ATP hydrolyzing enzymes such as transport ATPases the turnover rate of SMC complexes is rather low. The *B. subtilis* SMC and the *E. coli* MukB proteins have a basal activity of around 0.3 s^-1^ and lower than 0.1 s^-1^ ATP per SMC complex (48). In both cases addition of the kleisin subunit stimulates the activity between 2-10 fold. The observed activity of the *C. glutamicum* MksB is even lower (0.01 s^-1^). Stimulation of this low basal activity by addition of the kleisin subunit MksF increased the activity also about 2 fold. Unlike other tested SMC complexes, addition of the kite subunit MksE further increases the basal activiy. Addition of DNA further stimulates ATPase activity of the complex up to 10 fold of the basal activity. Interestingly, addition of the individual subunits MksF and MksG led to a decrease of the activity, while assembly of the whole MksBEF complex leads to highest ATPase activities in the presence of plasmid DNA. Addition of the MksG subunit decreases the activity of the DNA stimulated complex, likely because MksG degrades the plasmid DNA in the course of the experiment. Thus, there are distinct differences in the regulation of the ATPase cycle between MksBEFG complex and other SMC complexes (39,49). ATP hydrolysis in condensin complexes is required for DNA loop extrusion and hence these complexes can be seen as molecular motors that act in consecutive steps on DNA. At this stage, the exact mechanims of how MksBEFG identifies and binds to plasmid DNA remains unclear, however, it seems logical that the MksBEFG complex senses DNA topology that is different in plasmid DNA compared to chromosomal DNA. The on-off rate of the complex to its substrate DNA might be much higher and thus the ATPase is not needed to induce repetitive cycles to power loop extrusion.

Here, we show that MksG is a nuclease that cleaves DNA in a divalent cation-dependent manner. While *in vitro* manganese ions seem to allow highest activity, *in vivo* magnesium ions are likely the cofactor for MksG activity. However, the distantly related gyrase from *Staphylococcus aureus* is also a manganese dependet enzyme (50) and hence, we cannot rule out that MksG may indeed use Mn^2+^ *in vivo*. MksG shares the TOPRIM fold of Topo VI. Mutational analysis, as well as the structure of MksG reveal the conserved ion binding site in MksG and TopoVIA proteins, confirming earlier bioinformatic predictions (4). The acidic residues that coordinate the ion can be superimposed well with the homologous residues from the *M. jannaschii* topo VI enzyme (26). The catalytically active tyrosine in MksG is likely Y258, positioned right above the ion binding site. The N-terminal domain of MksG might act as a DNA clamp, helping to position or recognize the plasmid DNA. A large patch of positively chared amino acids is located at the inner face of the tip of the N-terminal domain, making this an ideal candiate for the DNA binding site. This places the MksG N-terminus in roughly the same position as the transducer domain of the TopoVIB subunit. The transducer in Topo VI connects the Bergerat ATPase domain with the TopoVIA subunit. The ATPase activity in Topo VI is required for DNA strand crossing. Strand crossing is likely not required in plasmid defense by MksG and hence this module might be superfluous. However, at this stage we can only speculate about the function of the condensin-like subunits MksBEF. ATP hydrolysis in MksB is not required in vitro for MksG activity, but presence of MksB in vivo has a clear effect on MksG dynamics and MksB is required for the plasmid defense activity as judged by the null allele phenotype. In vitro the nuclease activity of *C. glutamicum* MksG was not dependent on the condensin subunits MksBEF. However, the activity of MksG in vitro is slow compared to other nucleases. Therefore, in vivo a regulatory effect of MkeBEF on MksG might occur.

Interaction of SMC proteins and topoisomerases is common. The direct interaction of a Topo IV and MukB is well established (29,51,52). Thus, the interaction of MksBEF and MksG may not come as a surprise. However, there seems to be a large difference in the way the topoisomerase-like proteins bind to the SMC complexes. In MukBEF systems topoisomerase IV binds to MukB (27,28), while we identified here the kleisin subunit as interaction partner of MksG. Since the structure of the entire complex is unknown the precise interactions of the individual subunits remain unclear. However, bacterial two hybrid data (13) and the biolayer interferometry data presented here, suggest that MksG is localized rather to the region of the kleisin ring. For the SMC complex, it has recently been suggested that the SMC/ScpAB complex contains two DNA binding sites (double chamber) with the so termed meta-chamber in the inter-head/kleisin/kite space (40,53). The interaction of MksG within this region might position it perfectly to execute its nuclease function on plasmid DNA (Fig. 8).

**Figure 8.**
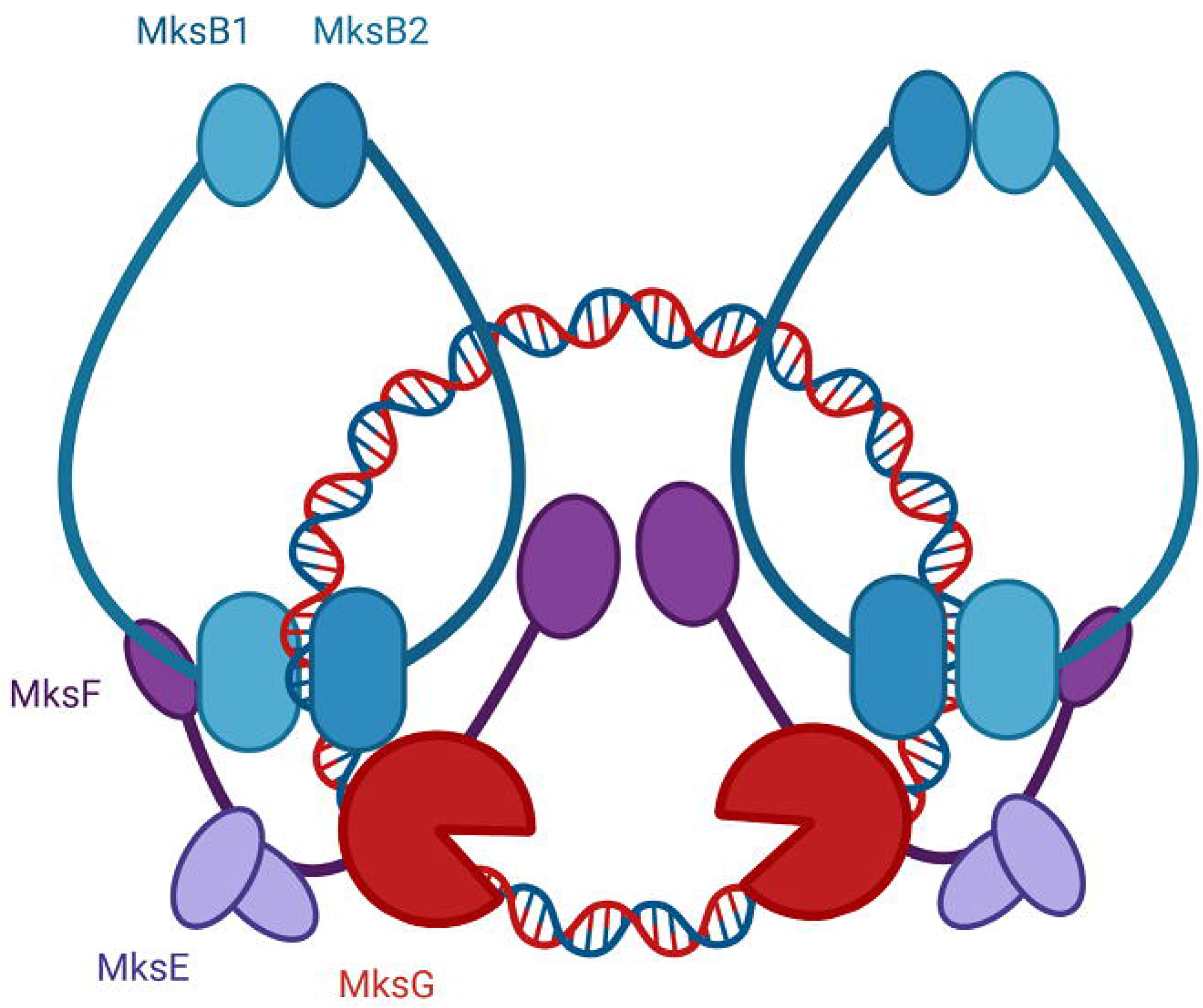
Model of the MksBEFG plasmid degradation system. The MukBEF homolog MksBEF likely forms a dimer of dimers that is linking the novel nuclease MksG via the kleisin subunit MksF. This complex organization positions the nuclease subunit in the vicinity of the DNA substrate that can be efficiently cleaved.

A fascinating aspect of the corynebacterial MksBEFG plasmid defense system is its spatial organization. We showed that all subunits localize to the cell poles in *C. glutamicum.* This polar localization seems to be dependent on the scaffold protein DivIVA. We have shown earlier that MksG and MksF interact with DivIVA in a bacterial two hybrid screen (13). We now show that depletion of DivIVA abolishes Mks complex localization, thereby further supporting the notion that likely a direct interaction between the Mks complex and DivIVA exists. DivIVA interacts with ParB in *C. glutamicum* and therefore localizes the *oriC* to the cell pole (54). However, the bulk of the chromosomal DNA is localized to the cell center. It remains to be tested where plasmids localize in *C. glutamicum,* but experiments with plasmid bearing strains lacking *mksB* reveal that there seems to be a polar enrichment. It is therefore attractive to hypothezize that the MksBEFG complex is polar localized and activates MksG nuclease activity there. Our single molecule tracking data suggest that MksG is less confined when plasmid DNA is transformed into the cells. This would be in line with a release of MksG from is polar scaffold. We hypothesize that, in absence of plasmids, MksG is held in standby at the cell poles by MksBEF. Upon binding to a plasmid, the complex loads and activates MksG that is then in turn released from the complex with the degraded plasmid. This spatial confinement might prevent toxic activation of the MksBEFG defense system elsewhere in the cell and erroneous nuclease activity on the chromosomal DNA.

## Data availability

Atomic coordinates and structure factors have been deposited in the protein data bank under the accession code 8B7F

## Acknowledgments

We gratefully acknowledge the core facilities PFBMI and PFC at the Institut Pasteur C2RT, as well as the synchrotron source Soleil (Saint-Aubin, France) for granting access to the facility and the staff of Proxima 1 for helpful assistance during X-ray data collection. The authors thank Dr. Kati Böhm for strains.

This work was funded by institutional grants from the Institut Pasteur, the CNRS, and Université Paris Cité. We are thankful to the Deutsche Forschungsgemeinschaft (DFG) for generous funding (BR 2915/6-2) to M.B.

## Material and Methods

### Bacterial strains, plasmids and oligonucleotides

#### Cloning

Strains, plasmids, and primers used in this study are listed in Supplementary Data Tables S2-4. Correct plasmid construction was controlled by DNA sequencing (Eurofins). *E. coli* NEB5α or *E. coli* NEB Turbo were used for cloning plasmids.

For heterologous protein expression, genes of interest (cg3103-cg3106) were amplified using PCR, digested with respective enzymes and ligated into pET28a (+) vector. Genes *mksF* (cg3106) *mksE* (cg3105) *mksG* (cg3103) were amplified using oligonucleotides MG045/MG046, MG047/MG048 and MG051/MG052, respectively, from genomic DNA and resulting amplicons were digested with NdeI/BamHI-HF. Digested fragments were ligated into digested pET28a-vector resulting in plasmids pMG001, pMG002, pMG003. Vector pET16b and gene *mksB* (cg3103) were amplified using primer pair MG093/MG094 and MG095/MG096 respectively and cloned using the Gibson Assembly Mastermix (NEB), resulting in plasmid pMG004. Primers were designed using https://nebuilder.neb.com/. For the MksB hydrolysis mutant (MksB^E1042Q^) a site-directed mutagenesis (Q5 site-directed mutagenesis Kit, NEB) was performed using primer pair MG060/MG061 and pMG004 as template, resulting in plasmid pMG005.

In order to generate the MksG mutants (MksG^E236A^, MksG^D279A^, MksG^Y258A^, MksG^Y276A^) site-directed mutagenesis (Q5 site-directed mutagenesis Kit, NEB) was performed, using primer pairs MG127/MG128, MG129/MG130, MG139/MG140 and MG141/MG142 respectively, plasmid pMG003 was used as template resulting in plasmids pMG006, pMG007, pMG017 and pMG018.

Derivatives of the suicide integration vector pK19mobsacB were used for clean knock-outs and allelic replacements in *C. glutamicum,* containing the modified genomic region interest including its 500bp up- and downstream homologous flanking sequences. In order to construct pMG009 primer pairs MG064/MG065 and MG066/MG067 were used to amplify the 500bp up- and downstream regions. The two fragments were used as a template for an overhang-PCR yielding in an 1000bp fragment, which was then digested with HindIII-HF and EcoRI-HF and subsequently ligated into digested pK19mobsacB vector. To construct pMG010 primer pairs MG080/MG081 and MG082/MG083 were used to amplify the 500bp up- and downstream regions. The two fragments were used as a template for an overhang-PCR yielding in an 1000bp fragment, which was then digested with XbaI and EcoRI-HF and subsequently ligated into digested pK19mobsacB vector. In order to construct pMG011 primer pairs MG084/MG085 and MG086/MG087 were used to amplify the 500bp up- and downstream regions. The two fragments were used as a template for an overhang-PCR yielding in an 1000bp fragment, which was then digested with HindIII-HF and EcoRI-HF and subsequently ligated into digested pK19mobsacB vector.

To construct a C-terminal tagged fusion proteins with Halo-Tag for an allelic replacement in *C. glutamicum* the Gibson Assembly technique was used. Primers were designed using https://nebuilder.neb.com/. Four fragments for each plasmid were amplified to generate plasmids pMG012-15. To amplify the pK19mobsacB backbone the primer pair MG097/MG098 were used for all plasmid derivatives (fragment 1). As a template for the Halo-Tag gene plasmid pK19mobsacB *parB*-Halo (Dr. G. Giacomelli, laboratory stock) was used. For plasmid pMG012 further three fragments were generated using primer pairs MG099/MG100, MG101/MG102 and MG103/MG104. For plasmid pMG013 further three fragments were generated using primer pairs MG105/MG106, MG107/MG108 and MG109/MG110. For plasmid pMG014 further three fragments were generated using primer pairs MG111/MG112, MG113/MG114 and MG115/MG116. For plasmid pMG015 further three fragments were generated using primer pairs MG117/MG118, MG119/MG120 and MG121/MG122.

Plasmid pMG016 was constructed by amplifying the *mksG* gene using primer pair MG089/MG090. Generated fragment was digested with HindIII and SalI and ligated into pXMJ19 vector.

#### Strain construction

Strains EMG007, EMG008, EMG009, EMG049, EMG051; and EMG053 were obtained by transforming plasmids pMG002, pMG001, pMG003, pMG006, pMG007 and pMG008 into chemical competent *E. coli* BL21 (DE3) pLysS cells. Strains EMG035 and EMG041 were obtained by transforming plasmids pMG004 and pMG005 into chemical competent *E. coli* Rosetta (DE3) pLysS cells.

Vectors were transformed via electroporation into *C. glutamicum* MB001 cells (55). Genomic integration of pK19mobsacB plasmids were selected on kanamycin, whereas the second crossover event was confirmed by growth on 10% sucrose. Screening of allelic replacements in *C. glutamicum* was confirmed by colony PCR. Strains CMG004, CMG005, CMG011 were generated by electroporation plasmids pMG009, pMG010 and pK19mobsacB *ΔmksB* into strain MB001. In order to obtain the strain *ΔmksG,* plasmid pMG011 was first electroporated and integrated into the MB001 genome. Secondly, plasmid pMG016 was transformed into the same strain. MksG expression from pXMJ19 plasmid was induced by 1mM IPTG. Once the second cross-over was confirmed (resulting in Δ*mksG*) by colony PCR, the cells were further grown in BHI and selected until plasmid pMG016 was lost again, resulting in strain CMG006.

To obtain strains CMG012, and CMG018 plasmid pMG012 was transformed into MB001 and CMG011. To obtain strains CMG013, CMG014 and CMG015 plasmids pMG013, pMG014 and pMG015 were transformed into MB001 cells.

Plasmid pBHK18 was transformed into MB001, CMG011, CMG012, CMG018, CMG004, CMG006 to generate strains CMG007, CMG023, CMG032, CMG034, CMG041 and CMG046, respectively. Plasmid pJC1 was transformed into MB001, CMG012, CMG018, CMG011, CMG004, CMG006 to generate strains CMG010, CMG033, CMG035, CMG038, CMG042 and CMG046, respectively.

The pSG-dCas9_sgRNA-*divIVA* plasmid was transformed into strain CBK114 to obtain CPF009.

For recombinant protein production in *E. coli* MksG was amplified from pMG003 using primers P1 and P2 and subcloned by Gibson Assembly into a pET28 derivative vector containing an N-terminal 6xHis-SUMO tag using primers P3 and P4. All plasmids were verified by Sanger sequencing (Eurofins Genomics, France).

### DivIVA depletion using the CRISPRi system

DivIVA was depleted as described by Giacomelli et al. (42).

### Plasmid Extraction from *C. glutamicum*

Plasmid extraction was performed as described by Böhm et al. (13).

### Genomic DNA Extraction

*C. glutamicum* cells were grown in 10ml BHI medium to exponential growth phase in presence of selection antibiotic (where necessary). Samples were adjusted to an OD_600_ of 16 per ml. Cell pellets were frozen and stored at −20°C overnight. DNA extraction was performed with Microbial DNA kit (Macherey Nagel), according to the manufacturer’s instructions.

### Real-Time PCR

DNA amplification was performed using 2x qPCR Mastermix SYBR blue (GeneON, Y220) according to the manufacturer’s manual, where reaction volumes of 25μl contained 200nM oligonucleotides and 10μl of diluted DNA, respectively. Samples were measured in technical triplicates via an AriaMx and Ct values were determined using the Agilent Aria Software 1.8v. Primer efficiencies were estimated by calibration dilution curves and slope calculation; data were analysed by 2^-ΔCT^ method accounting for dilution factors and sample volumes used for DNA purification. Same primers were used as mentioned by Böhm et al. (13).

### Protein Purification

Heterologous expression of His_6_-MksE and His_6_-MksG was performed in *E. coli* strains EMG007 and EMG009, respectively, in LB-medium containing the appropriate antibiotics [Kanamycin 50 μg ml^-1^ Chloramphenicol 34 μg ml^-1^]. An IPTG concentration of 1mM was added at an optical density at 600nm of ~0.6 to induce protein expression. Cells were grown for 3-4h at 37°C.

Heterologous expression of His_6_-MksF, His10-MksB and His10-MksB^E1042Q^ was performed in *E. coli* strains EMG008, EMG035 and EMG041, respectively, in LB-medium containing the appropriate antibiotics [Kanamycin 50 μg ml^-1^ Chloramphenicol 34 μg ml^-1^]. An IPTG concentration of 0.5 mM was added at an optical density at 600nm of 0.6-0.9 to induce protein expression at 18°C. Cells were grown for ~16h.

The cultures were harvested at 6,500 rcf for 15 min at 4°C. Cell pellets were frozen and stored at −80°C until needed.

Cell pellets were resuspended in cold Buffer IMAC [50mM Tris-HCl pH 8.5, 0.5M NaCl, 10mM MgCl_2_, 10% glycerol] and supplemented with 15mM Imidazole, a pinch of lyophilized DNaseI and Protease Inhibitor Cocktail (cOmplete™, EDTA-free Protease Inhibitor Cocktail, Roche 04693132001). The homogenous suspension was passed through a French Pressure cell (Amico) in order to disrupt the cells. The solution was passed through three times at 20,000 psi inner cell pressure. The suspension was centrifuged at 10,000 rpm at 4°C in Beckman Coulter Avanti J-25 Centrifuge using the JA-10 rotor with falcon adapters to eliminate cell debris. The supernatant was transferred to a fresh falcon and directly used for immobilized metal affinity chromatography (IMAC) using the Äktapure25 system with a sample pump (Cytiva) and 1ml Ni-NTA columns (Macherey-Nagel). Proteins were concentrated where necessary using Amicon filter devices with appropriate molecular weight cut-off (Millipore; Merck).

For MksE, MksF, MksG, MksG^E236A^, MksG^D279A^, MksG^Y258A^ and MksG^Y276A^ purifications, size-exclusion chromatography followed using Superdex200 Increase 10/300 GL (Cytiva) and fractions were pooled and concentrated snap frozen and stored at −80°C or directly used for the assays. Purifying MksE and MksF together yielded in cleaner protein fractions. However, it was not possible to separate the two proteins via size-exclusion or anion-exchange chromatography.

For MksB and MksB^E1042Q^ purifications, fractions after IMAC were diluted in Buffer without NaCl to an end concentration of ~150 mM NaCl and chromatographed using Heparin HiTrap HP column (5ml) (Cytiva) and resulted in a sharp peak at 300mM NaCl. The fractions were pooled and concentrated and chromatographed with SEC Buffer [50mM Tris-HCl pH 8.5, 0.5 M NaCl, 10 mM MgCl_2_, 1 mM DTT, 5 % glycerol] using Superdex200Increase 10/300 GL or Superose6 Increase 10/300 GL (Cytiva). Peak fractions were pooled, concentrated, snap frozen and stored at −80°C or directly used for the assays. Protein purities were analyzed via SDS-PAGE using 8-12% polyacrylamide gels.

### Protein expression and purification for crystallization

N-terminal 6xHis-SUMO-tagged constructs were expressed in *E. coli* BL21 C41 following an autoinduction protocol. After 4h at 37 °C, cells were grown for 20 h at 20°C in 2YT complemented autoinduction medium (56) containing 50μg/ml Kanamycin. Cells were harvested and flash frozen in liquid nitrogen. Cell pellets were resuspended in lysis buffer (50 mM TRIS-HCl pH 8.0,5 mM NaCl, 5% glycerol, 1 mM MgCl_2_, benzonase, lysozyme, EDTA-free protease inhibitor cocktails (ROCHE)) at 4°C and lysed using a CF Disruptor cell disintegrator system (CellD.com). The lysate was centrifuged (13000 rpm) for 1 h at 4 °C, and the supernatant was loaded onto a Ni-NTA affinity chromatography column (HisTrap FF crude, Cytiva) pre-equilibrated in buffer A (50 mM TRIS-HCl pH 8.0, 500 mM NaCl, 5% glycerol, 10 mM imidazole). His-tagged proteins were eluted with a linear gradient of buffer B (50 mM TRIS-HCl pH 8.0, 500 mM NaCl, 5% glycerol, 1 M imidazole). The fractions of interest were pooled and dialyzed at 4°C overnight in SEC buffer (20 mM TRIS-HCl pH 9.0, 500 mM NaCl, 5% glycerol). The protein was concentrated and loaded onto a Superdex 200 16/60 size exclusion column (GE Healthcare) pre-equilibrated at 4 °C in SEC buffer. The peak corresponding to the protein was concentrated, flash frozen in liquid nitrogen and stored at −80°C. Purity was verified by sodium dodecyl sulfate–polyacrylamide gel electrophoresis (SDS-PAGE).

### Crystallization

Initial screening of crystallization conditions was carried out by the vapor diffusion method using a MosquitoTM nanoliter-dispensing system (*TTP Labtech, Melbourne, United Kingdom)* following established protocols (57). MksG has a high propensity to crystallize however the diffraction properties are very poor, very likely due to the flexibility between the C- and N-ter domains. The best crystals of MksG (20 mg/mL) were obtained after 3 days in 10% (w/v) PEG 6K, 30% (v/v) ethanol and 10 mM sodium acetate. Crystals were cryoprotected in crystallization solution supplemented by 30% (v/v) of ethylene glycol before flash freezing in liquid nitrogen.

### Data collection, structure determination, and refinement

X-ray diffraction data were collected at 100 K at the PROXIMA 1 beamline of the SOLEIL Synchrotron (Saint-Aubin, France). The dataset was processed using XDS (58) and AIMLESS from the CCP4 suite (59) (see Table S1). Merged data was further subjected to anisotropy correction with STARANISO (60), the elliptical diffraction limits were dh00 = 5.3 Å, d0k0 = 4.3 Å, d00l = 4.3 Å. The structure was solved by molecular replacement methods, using Phaser from the CCP4 suite, with the individual N- and C-terminal domains generated by Alphafold2 as search models. The structures were refined through iterative cycles of manual model building with COOT (61) and reciprocal space refinement with PHENIX (62) or BUSTER (63). Non-crystallographic symmetry and secondary structure restraints were applied. The final refinement statistics are shown in Table S1 and a representative view of the final electron density map is shown in Supplementary Figure 1B. Structural figures were generated with ChimeraX (64).

### AlphaFold prediction and structure-based analysis

A structural model of MksG was calculated using AlphaFold2 (65) installed on local servers (Supplementary Figure 1A). The structural homology search was performed against the Alphafold database on a local installation of DaliLite (66).

### Electrophoretic mobility shift assay

DNA-MksB/ DNA-MksG/ DNA-MksEF binding was assayed by using purified protein and circular plasmid DNA (pJC1 or pBHK18), PCR-amplified (pK19mobsacB) linear double-stranded DNA or single-stranded DNA (49bp oligonucleotide, MG105). 0 - 10μg of protein were incubated with 200ng of linear or circular plasmid DNA or 30μM ssDNA for 30min@ 30°C in Buffer: 20mM Tris-HCl pH8.5, 50mM NaCl, 10mM MgCl_2_. Samples were separated on 0.7% agarose gels at 4V/cm for 2.5h (with 5x Nucleic buffer 50mM Tris-HCl pH 8.5, 25% glycerol, 5mM EDTA, 0.2% bromophenol blue, 0.2% xylene cyanole FF) in TAE Buffer. After the separation, agarose gels were stained in GelRed or Ethidium Bromide.

### Cleavage Assays

Nicking Assays were performed as described by Xiong et al. (67). The reaction contained 30ng/μl plasmid DNA (extracted from C.gl *ΔmksB* strain (CMG011)) in 50mM potassium acetate, 20mM Tris-acetate (pH7.9), 10mM metal ions/ EDTA, 0.1mg ml^-1^ BSA and 10μM MksG. The reactions were incubated at 30°C and samples were taken after 0.5, 1 and 3h. The reaction was stopped by adding 6x Purple DNA-Loading dye (NEB) and boiled at 90°C for 5min. Samples were stored at −20°C until run on a 0.7-1% agarose gels in TAE at 6V/cm for 1.5h. DNA was post-stained with GelRed or Ethidiumbromide.

### ATP hydrolysis assays

ATP hydrolysis was analysed in steady state reactions using the EnzChek Phosphate Assay Kit (E-6646, Molecular Probes Inc.) according to the manufacturer’s manual. Reaction volumes were reduced to 100μl and assayed in flat-bottom 96-well plates (Greiner-UV-Star 96-well plates). Reactions were measured at 30°C over a time-course of 3 h in a Tecan Infinite 200 Pro with the Software Tecan i-control v.2.0. The reactions contained 4μM of MksB or MksB^E1042Q^ as well as the additional subunits (unless otherwise stated), 2mM Mg-ATP (unless otherwise stated) and 50ng/μl plasmid DNA (pBHK18) where indicated. Before starting the reactions with 2mM Mg-ATP, the reactions were preincubated for 15min in order to eliminate phosphate contamination. Data was analysed with Graph Pad Prism 5.

### Bio-Layer Interferometry

Measurements were performed on the BLItz platform (FortéBio (now Sartorius)) using Streptavidin (SA) Biosensors (Sartorius, 18-5019). The BLItz device was operated with the

Software BLItz Pro (version 1.3.1.3) using the Advanced Kinetics Protocol modified as shown in **Fehler! Verweisquelle konnte nicht gefunden werden.**.

Beforehand, bait proteins were biotinylated (ratio 3:1) using the EZ-Link NHS-PEG4 Biotinylation kit (ThermoScientific, 21455) as recommended by the manufacturer. Buffer exchange was performed using Zeba Spin Desalting Columns 7k MWCO 5 ml (ThermoScientific; 89891).

Biosensors were hydrated for at least 10 min in a 96-well plate in 200μl BLI-Buffer [10 mM Na_2_HPO_4_, 137 mM NaCl, 2.7 mM KCl, 1.8 mM KH_2_PO_4_, 10 mM MgCl_2_, 5 % Glycerol, pH 7.4].

As a starting point, unspecific binding was tested. For this, unbiotinylated protein was used to be loaded onto the Streptavidin biosensor. Additionally, loaded biotinylated-protein was tested with BLI Buffer only in order to rule out unspecific binding of the buffer. For the Binding assays the initial baseline was measured in a black 0.5 ml reaction tube filled with 250μl BLI Buffer. Next, 4μl of bait protein was pipetted into the drop holder of the machine and loaded onto the biosensor. Another baseline was measured in the same tube as before. For the association the analyte protein (in different concentrations) was pipetted into the cleaned drop holder. For the dissociation, the machine was shifted back to the tube position. For each measurement a fresh biosensor was used.

Kinetic analysis was performed by the Software BLItz Pro (version 1.3.1.3).

### Microscopic analyses

#### Fluorescence microscopy

*C. glutamicum* cells were grown in BHI medium (OXOID) at 30 °C, 200 rpm. Plates and liquid cultures were supplement with 50 μg mL*-*1 kanamycin when needed (CMG012, CMG013, CMG014, CMG015). Colonies were picked from freshly plated stocks and grown overnight. The next morning, a fresh culture with an OD_600_ of 0.5 was started and grown until an OD_600_ of ~2. 1mL of the culture was stained with 250nM HaloTag TMR Ligand (Promega) for 30 min at 30 °C. Cells were harvested (4000 rpm, 3 min, 30 *°*C) and washed five times in PBS Buffer, pH 7.4 (sterile filtered 0.2 μm).

Image Analysis and Data processing was performed with FIJI Software’s (68) and Plug-in MircobeJ (69).

#### Single Molecule Localization Microscopy/ Single Particle Tracking

*C. glutamicum* cells were grown in BHI medium (OXOID) at 30 °C, 200 rpm. Plates and liquid cultures were supplement with 50 μg mL*-*1 kanamycin when needed (CMG015, CMG012, CMG018, CMG032, CMG034). Colonies were picked from freshly plated stocks and grown overnight. The next morning, a fresh culture with an OD_600_ of 0.5 was started and grown until an OD_600_ of ~2. 1mL of the culture was stained with 50nM HaloTag TMR Ligand (Promega) for 30 min at 30 °C. Cells were harvested (4000 rpm, 3 min, 30 *°*C) and washed five times in TSEMS [50mM Tris, pH 7.4, 50mM NaCl, 10mM EDTA, 0.5 % sucrose (sterile filtered 0.2 μm)] osmoprotective buffer.

Slides were cleaned with 1 M KOH, rinsed with ddH2O and dried with pressurized air. Agar pads where prepared with the aid of gene frames (Thermo Fisher, Dreieich, Germany) and 1% (w/v) low melting agarose (agarose, low gelling temperature, Sigma-Aldrich, Taufkirchen, Germany) in sterile filtered TSEMS (0.2 μm pores)..

SMLM imaging was performed with an Elyra 7 (Zeiss) inverted microscope equipped with two pco.edge sCMOS 4.2 CL HS cameras (PCO AG), connected through a DuoLink (Zeiss), only one of which was used in this study. Cells were observed through an alpha Plan-Apochromat 63×/1.46 Oil Korr M27 Var2 objective in combination with an Optovar 1 × (Zeiss) magnification changer, yielding a pixel size of 97 nm. During image acquisition, the focus was maintained with the help of a Definite Focus.2 system (Zeiss). Fluorescence was excited with a 561 nm (100 mW) laser, and signals were observed through a multiple beam splitter (405/488/561/641 nm) and laser block filters (405/488/561/641 nm) followed by a Duolink SR QUAD (Zeiss) filter module (secondary beam splitter: LP 560, emission filters: EF BP420-480 + BP495-550).

For each time lapse series, 10,000 frames were taken with 20 ms exposure time (~24 ms with transfer time included) and 50% 561 nm intensity laser in TIRF mode (62° angle).

For single-particle tracking, spots were identified with the LoG Detector of TrackMate v6.0.1 (70), implemented in Fiji 1.53 g (71), an estimated diameter of 0.5 μm, and median filter and sub-pixel localization activated. The signal to noise threshold for the identification of the spots was set at 5. To limit the detection of ambiguous signal, frames belonging to the TMR bleaching phase (first 500 frames) were removed from the time lapses prior to the identification of spots. Spots were merged into tracks via the Simple LAP Tracker of TrackMate, with a maximum linking distance of 500 nm, two frame gaps allowed, and a gap closing max distance of 800 nm. Only tracks with a minimum length of 5 frames were used for further analysis, yielding a minimum number of total tracks per sample of 1132.

To identify differences in protein mobility and/or behavior, the resulting tracks were subjected to dwell time, mean-squared-displacement (MSD), and square displacement (SQD) analysis in SMTracker 2.0 (72), described previously (73).

Dwell time distribution was determined for a confinement radius of 97 nm and fitted with two components. The average MSD was calculated for four separate time points per strain (exposure of 20 ms—τ = 24, 48, 72, and 96 ms), followed by fitting of the data to a linear equation. The last time point of each track was excluded to avoid track-ending-related artifacts. The cumulative probability distribution of the square displacements (SQD) was used to estimate the diffusion constants and relative fractions of up to three diffusive states. Diffusion constants were determined simultaneously for the compared conditions (MksG and related mutants), therefore allowing for a more direct population fraction comparison.

#### Statistics

Mean square displacement, jump distance, dwelling time analysis and related statistics were performed via SMTracker 2.0. (72) FRAP-related statistics were performed via Rstudio (R studio version 1.4.1106, R version 4.0.5 (31 March 2021) (74). The following packages were used: ggplot2 (75), PMCMRplus (76), and nlstools (77).

