## Supplemental Material for "A novel nuclease is the executing part of a bacterial plasmid defense system"

Running title: The MksBEFG plasmid defense system

[#] These authors contributed equally

* To whom correspondence should be addressed: Marc Bramkamp, Christian-Albrechts-University Kiel, Institute for General Microbiology, Am Botanischen Garten 1-9, 24118 Kiel, Germany,; Phone: +49 (0)431-880-4341; Telefax: +49(0)431-880-2198, Twitter: @BramkampLab

**Table S1. Crystallographic data collection and refinement statistics.**

| **Data collection** | **MksG** |
|---|---|
| Synchrotron Beamline | SOLEIL Proxima 1 |
| Wavelength (Å) | 0.9786 |
| Space group | $P2_12_12_1$ |
| Cell dimensions *a*, *b*, *c* (Å) | 78.086, 126.776, 150.111 |
| Resolution (Å) | 61.5 – 4.6 (4.77 – 4.6)* |
| *R*pim | 0.054 (0.444) |
| *I* / *s*(*I*) | 9.6 (1.6) |
| Completeness (%) | 99.5 (94.1) |
| CC(1/2) | 0.998 (0.816) |
| Multiplicity | 13.1 (12.7) |
| Total observations | 114710 |
| Unique observations | 8738 (597) |
| **Refinement** | |
| Resolution (Å) | 4.6 |
| No. reflections | 7594 ** |
| *R*work/ *R*free (%) | 0.27 / 0.311 |
| No. atoms | |
| Protein | 5862 |
| Ligands/ions | - |
| Solvent | - |
| Average B-factors ($Å^2$) | |
| Protein | 147 |
| Ligand/ions | - |
| Solvent | - |
| R.m.s deviations | |

| | |
|---|---|
| Bond lengths (Å) | 0.004 |
| Bond angles (º) | 0.888 |
| Ramachandran favored (%) | 97.4 |
| Ramachandran outliers (%) | 0.27 |
| **PDB code** | 8B7F |

*Values in parenthesis correspond to the highest resolution shell.

**Number of unique reflections used for structure determination and refinement, after the merged dataset was subjected to anisotropic resolution surface cut-off with StarAniso (see Methods).

**Table S2: Strains used in this study**.

| Strain Name | Description | Genotype | Reference |
|---|---|---|---|
| | *Escherichia coli* | | |
| BL21 (DE3) pLysS | | F– dcm ompT hsdS(rB – mB – ) gal λ(DE3) [pLysS CamR ] | Novagen |
| EMG007 | BL21 (DE3) pLysS/ pET28a_His6-MksE | BL21 (DE3) pLysS derivative, pET28a mksE | This study |
| EMG008 | BL21 (DE3) pLysS/ pET28a_His6-MksF | BL21 (DE3) pLysS derivative, pET28a mksF | This study |
| EMG009 | BL21 (DE3) pLysS/ pET28a_His6-MksG | BL21 (DE3) pLysS derivative, pET28a mksG | This study |
| EMG049 | BL21 (DE3) pLysS/ pET28a_*mksG*$^{D279A}$ | BL21 (DE3) pLysS derivative, pET28a mksG$^{D279A}$ | This study |
| EMG051 | BL21 (DE3) pLysS *pET28a_mksG*$^{E236A}$ | BL21 (DE3) pLysS derivative, pET28a mksG$^{E236A}$ | This study |
| EMG055 | *E. coli* BL21 (DE3) pLysS/ pET28a_*mksG*$^{Y258A}$ | BL21 (DE3) pLysS derivative, pET28a mksG$^{Y258A}$ | This study |
| EMG057 | *E. coli* BL21 (DE3) pLysS *pET28a_mksG*$^{Y276A}$ | BL21 (DE3) pLysS derivative, pET28a mksG$^{Y276A}$ | This study |
| BL21 C41 | BL21 C41 expression strain | F – ompT hsdSB (rB- mB-) gal dcm (DE3) | Sigma Aldrich |
| Rosetta (DE3)pLysS | | F- ompT hsdSB(rB- mB-) gal dcm (DE3) pLysSRARE (CamR) | Novagen |
| EMG035 | Rosetta (DE3) pLysS pET16b-*mksB* | Rosetta (DE3) pLysS derivative, pET16b mksB | This study |
| EMG041 | Rosetta pLysSRARE pET16b_mksB $^{E1042Q}$ | Rosetta (DE3) pLysS derivative, pET16b mksB$^{E1042Q}$ | This study |
| | *Corynebacterium glutamicum* | | |
| MB001 | MB001 | ATCC 13032 with in-frame deletion of prophages CGP1 (cg1507-cg1524), CGP2 (cg1746-cg1752), and CGP3 (cg1890-cg2071) | Baumgart et al. (1) |
| CMG004 | MB001 *ΔmksF* | MB001 derivative, *ΔmksF* | This study |
| CMG005 | MB001 *ΔmksE* | MB001 derivative, *ΔmksE* | This study |
| CMG006 | MB001 *ΔmksG* | MB001 derivative, *ΔmksG* | This study |
| CMG007 | MB001 pBHK18 | MB001 derivative, pBHK18 | This study |
| CMG010 | MB001 pJC1 | MB001 derivative, pJC1 | This study |
| CMG011 | MB001 *ΔmksB* | MB001 derivative, *ΔmksB* | This study |
| CMG012 | MB001 *mksG::mksG-halo-tag* | MB001 derivative, *mksG::mksG-halo-tag* | This study |
| CMG013 | MB001 *mksE::mksE-halo-tag* | MB001 derivative, *mksE::mksE-halo-tag* | This study |
| CMG014 | MB001 *mksF::mksF-halo-tag* | MB001 derivative, *mksF::mksF-halo-tag* | This study |
| CMG015 | MB001 *mksB::mksB-halo-tag* | MB001 derivative, *mksB::mksB-halo-tag* | This study |
| CMG018 | MB001 *ΔmksB mksG::mksG-halo-tag* | MB001 derivative, *ΔmksB mksG::mksG-halo-tag* | This study |
| CMG023 | CMG011 + pBHK18 | MB001 derivative, *ΔmksB* pBHK18 | This study |

| | | | |
|---|---|---|---|
| CMG032 | CMG012 + pBHK18 | MB001 derivative, *mksG::mksG-halo-tag* | This study |
| CMG033 | CMG012 + pJC1 | MB001 derivative, *mksG::mksG-halo-tag* | This study |
| CMG034 | CMG018 + pBHK18 | MB001 derivative, ΔmksB *mksG::mksG-halo-tag* | This study |
| CMG035 | CMG018 + pJC1 | MB001 derivative, *ΔmksB mksG::mksG-halo-tag* | This study |
| CMG038 | CMG011 + pJC1 | MB001 derivative, *ΔmksB* pJC1 | This study |
| CMG041 | CMG004 + pBHK18 | MB001 derivative, *ΔmksF* pBHK18 | This study |
| CMG042 | CMG004 + pJC1 | MB001 derivative, *ΔmksF* pJC1 | This study |
| CMG045 | CMG006 + pBHK18 | MB001 derivative, *ΔmksG* pBHK18 | This study |
| CMG046 | CMG006 + pJC1 | MB001 derivative, *ΔmksG* pJC1 | This study |
| RES167 | RES167 *(Restriction-deficient mutant, otherwise considered wild-type)* | | Tauch et al. (2) |
| CBK114 | RES167 *mksB::mksB-mNeonGreen* | RES 167 derivative, *mksB::mksB-mNeonGreen* | Lab collection |
| CPF009 | RES167 *mksB::mksB-mNeonGreen*, dCas-*divIVA* | RES 167 derivative, *mksB::mksB-mNeonGreen*, dCas-*divIVA,* pSG-dcas9_sgRNA-*divIVA* (IPTG-inducible) | This study |

**Table S3: Plasmids used in this study**.

| **Name** | **Description** | **Reference** |
|---|---|---|
| pET28a(+) | *E. coli* expression vector, carrying an N-terminal 6xHisTag/thrombin/T7Tag configuration plus an optional C-terminal HisTag sequence, Kan$^r$ | Novagen |
| pMG001 | pET28a(+) *mksF* | This study |
| pMG002 | pET28a(+) *mksE* | This study |
| pMG003 | pET28a(+) *mksG* | This study |
| pMG006 | pET28a_*mksG*$^{D279A}$ | This study |
| pMG007 | *pET28a_mksG*$^{E236A}$ | This study |
| pMG008 | *pET28a_mksG*$^{E236A, D279A}$ | This study |
| pMG017 | *pET28a_mksG*$^{Y258A}$ | This study |
| pMG018 | *pET28a_mksG*$^{Y276A}$ | This study |
| pMBA001 | pET28a SUMO-*mksG* | This study |
| pET16b | *E. coli* expression vector, carrying an N-terminal His•Tag® sequence followed by a Factor Xa site and three cloning sites, *bla* | Novagen |
| pMG004 | pET16b-*mksB* | This study |
| pMG005 | pET16b_*mksB*$^{E1042Q}$ | This study |
| pK19*mobsacB* | Kan$^r$; plasmid for allelic exchange in C. glutamicum (pK18 *oriV*$_{E.c.}$ *sacB lacZ*α) | Schäfer et al. (3) |
| pK19msB-Δ*mksB* | Plasmid to generate a *mksB* deletion, pK19msB-Δ*mksB* | Böhm et al. (4) |
| pMG009 | pK19 *mobsacB ΔmksF* | This study |

| | | |
|---|---|---|
| pMG010 | pK19 *mobsacB ΔmksE* | This study |
| pMG011 | pK19 *mobsacB ΔmksG* | This study |
| pMG012 | pK19 *mobsacB -mksG-halo* | This study |
| pMG013 | pK19 *mobsacB -mksE-halo* | This study |
| pMG014 | pK19 *mobsacB -mksF-halo* | This study |
| pMG015 | pK19 *mobsacB -mksB-halo* | This study |
| pJC1 | *E. coli/ C. glutamicum* shuttle vector, $Kan^r$ | Cremer et al. (5) |
| pBHK18 | Based on pNG2 plasmid*, aph(3' ($Kan^r$)-IIa* | Kirchner and Tauch (6) |
| pXMJ19 | *E. coli/ C. glutamicum* shuttle vector, p*tac*,*$lacI^q$*, $Cam^r$ | Jakoby et al. (7) |
| pMG016 | pXMJ19 *mksG* | This study |
| pSG-dCas9_sgRNA-*divIVA* | Ptac, $lacI^q$, repA, ori PUC, dCas9, *diVIVA*-sgRNA,$Kan^r$ | Giacomelli et al. (8) |

**Table S4: Oligonucleotides used in this study. Specific restriction sites are underlined.**

| Name | Description | Sequence 5' - 3' | Restriction site |
|---|---|---|---|
| MG045 | NdeI_MksF_F | TAATGCATATGACCGTTGTATCG | NdeI |
| MG046 | MksF_BamHI_R | TACTGGATCCTCATTTATCCATCTC | BamHI |
| MG047 | NdeI_MksE_F | GATAGCATATGAATGATCAGCTGT | NdeI |
| MG048 | MksE_BamHI_R | CTATGGATCCTCACTTCTGTTCC | BamHI |
| MG049 | NdeI_MksB_F | TAATGCATATGACCAGCGAACAAG | NdeI |
| MG050 | MksB_EagI_R | ATGACGGCCGTTATTTCTCGATCC | EagI |
| MG051 | NdeI_MksG_F | ATAGCATATGCCATTGTTTATCGAC | NdeI |
| MG052 | MksG_BamHI_R | CGATGGATCCTCACCCACGAATT | BamHI |
| MG053 | seq-MksB_middle1_F | AATTTTGAGTGCGAAGAGG | / |
| MG055 | seq-MksB_middle2_F | AAGAAATCGCGCGGGAAG | / |
| MG057 | seq-MksF_middle_F | CAGATTGAAGCGGTCCAC | / |
| MG058 | seq-MksB_middle_R1 | GTTGTTGAGCTCCAAAAGC | / |
| MG059 | seq-MksB_middle_R2 | CTCATTGGCATCGATTTG | / |
| MG060 | SDM_MksB_G->C_E1042Q_F | CATTCTGGACcAAGCCTTCGACCGC | / |
| MG061 | SDM_MksB_R | ACGGTGGCGTAGGTGGGA | / |
| MG064 | HindIII_500up_mksF-ko_F | CATAAGCTTCTCGTGGGAACCCGCCAA | HindIII |
| MG065 | 500up_mksF-ko_R | GGAATGGAGTATGGAAGTTGGGCCAATTGACTATTCCAG | / |
| MG066 | 500down_mksF-ko_F | CCAACTTCCATACTCCATTCCTTTGGCAGAAAGCGAGAT | / |
| MG067 | EcoRI_mksF_500down_R | CATGAATTCGTCTTTTTAGCTAATGAATAATCCA | EcoRI |
| MG080 | XbaI_500up_mksE_F | CATTCTAGAGCTTCGCGATACCCGCAG | XbaI |
| MG081 | 500up_mksE-ko_R | GGAATGGAGTATGGAAGTTGGCATTCATTTATCCATCTCGC | / |
| MG082 | mksE-ko_500down_F | CCAACTTCCATACTCCATTCCGGAAATGAAGAGGAACAGAA | / |
| MG083 | EcoRI_mksE_500down_R | CATGAATTCGTCTTGATCAACGGGAAACAC | EcoRI |
| MG084 | HindIII_500up_mksG_F | CATGAAGCTTACAAGCCTTCGCCCGTTATG | HindIII |
| MG085 | 500up_mksG-ko_R | GGAATGGAGTATGGAAGTTGGGTTATTTCTCGATCCTAGAGAA | / |
| MG086 | mksG-ko_500down_F | CCAACTTCCATACTCCATTCCGTTGAAGAAATCGAGAAAGT | / |
| MG087 | EcoRI_mksG_500down_R | CATGGAATTCGCGCCCTCCATATCGCA | EcoRI |
| MG088 | BamHI_MksG_F | CATATGGATCCATGCCATTGTTTATC | BamHI |
| MG089 | MksG_SalI_R | CATATGTCGACTCACCCACGAATTAC | SalI |
| MG090 | HindIII_mksG_F | CATATAAGCTTATGCCATTGTTTATC | HindIII |

| | | | |
|---|---|---|---|
| MG093 | pET16b_fwd | ctcgaggatccggctgcta | XhoI |
| MG094 | pET16b_rev | catatgacgaccttcgatatgg | NdeI |
| MG095 | mksB_fwd | atatcgaaggtcgtcatatggtgaccagcgaacaagctttag | NdeI |
| MG096 | mksB_rev | ttagcagccggatcctcgagttatttctcgatcctagagaaactg | (XhoI) |
| MG097 | pK19msB_fwd | GTCGACTCTAGAGGATCC | / |
| MG098 | pK19msB_rev | CTGCAGGCATGCAAGCTT | / |
| MG099 | mksG last 500 bp_fwd | tgattacgccaagcttgcatgcctgcagGACTTGGTGACGCCGAAG | / |
| MG100 | mksG last 500 bp_rev | atttccatacctgcCCCACGAATTACTTTCTCGATTTC | / |
| MG101 | halo-tag_fwd | aagtaattcgtgggGCAGGTATGGAAATCGGTAC | / |
| MG102 | halo-tag_rev | tcgaggcggatgcgTTAGGAAATCTCCAGAGTAGAC | / |
| MG103 | mksG 500 down_fwd | tggagatttcctaaCGCATCCGCCTCGATGTTGC | / |
| MG104 | mksG 500 down_rev | cggtacccggggatcctctagagtcgacCACCAGCGCCGCGCCCTC | / |
| MG105 | mksE last500bp_fwd | tgattacgccaagcttgcatgcctgcagTATCACCACAGATCAAGATGC | / |
| MG106 | mksE last500bp_rev | atttccatacctgcCTTCTGTTCCTCTTCATTTCC | / |
| MG107 | halo-tag_fwd | aagaggaacagaagGCAGGTATGGAAATCGGTAC | / |
| MG108 | halo-tag_rev | gctgctttgttaaaTTAGGAAATCTCCAGAGTAGAC | / |
| MG109 | mksE 500bp down_fwd | tggagatttcctaaTTTAACAAAGCAGCCCATG | / |
| MG110 | mksE 500bp down_rev | cggtacccggggatcctctagagtcgacCGCATTGATGTCTTGATCAAC | / |
| MG111 | mksF last500bp_fwd | tgattacgccaagcttgcatgcctgcagTCGCAGAGAGCCGACGCATG | / |
| MG112 | mksF last500bp_rev | atttccatacctgcTTTATCCATCTCGCTTTCTGCCAAATC | / |
| MG113 | halo-tag_fwd | gcgagatggataaaGCAGGTATGGAAATCGGTAC | / |
| MG114 | halo-tag_rev | tgtttaaacccctgTTAGGAAATCTCCAGAGTAGAC | / |
| MG115 | mksF 500bpdown_fwd | tggagatttcctaaCAGGGGTTTAAACAGTGAAG | / |
| MG116 | mksF 500bpdown_rev | cggtacccggggatcctctagagtcgacGGTGTCTCTGTCTTTTTAGC | / |
| MG117 | mksB last500bp_fwd | tgattacgccaagcttgcatgcctgcagACCAGCGGTGACCTGGGAAC | / |
| MG118 | mksB last500bp_rev | atttccatacctgcTTTCTCGATCCTAGAGAAACTGGAATTG | / |
| MG119 | halo-tag_fwd | ctaggatcgagaaaGCAGGTATGGAAATCGGTAC | / |
| MG120 | halo-tag_rev | ctagagaaactggaTTAGGAAATCTCCAGAGTAGAC | / |
| MG121 | mksB 500bp down_fwd | tggagatttcctaaTCCAGTTTCTCTAGGATCGAG | / |
| MG122 | mksB 500bp down_rev | cggtacccggggatcctctagagtcgacTGGTTTTCGAGCCATTTG | / |
| MG127 | SDM_MksG_A->C_E236A_F | CTGATGGTGGcAAACCTCGATTC | / |
| MG128 | SDM_MksG-EA_R | AATTACTTGCGGTTCTTG | / |
| MG129 | SDM_MksG_A->C_D279A_F | TACTGGGGTGcaCTTGACCTGG | / |
| MG130 | SDM_MksG-DA_R | CAGCAACCGACCATTAGA | / |
| MG139 | SDM_MksG_TAC->GCA_Y258A_F | GGGTGCGGGCgcaCGTGCAGTAG | / |
| MG140 | SDM_MksG-Y258A_R | CAAGCAATTGTTACGCCC | / |
| MG141 | SDM_MksG_TAC->GCA_Y276A_F | TCGGTTGCTGgcaTGGGGTGACC | / |
| MG142 | SDM_MksG-Y276A_R | CCATTAGAAAAGTAGGGTC | / |
| P1 | | GAGGCTCACCGCGAACAGATTGGTGGCATGCCATTGTTTATCGACGA | |
| P2 | | CTTCCTTTCGGGCTTTGTTAGCAGCCGGATCTTTACCCACGAATTACTTTCTC | |
| P3 | | AGATCCGGCTGCTAACAAAGCCCGAAAGGAAG | |
| P4 | | GCCACCAATCTGTTCGCGGTGAGCCTC | |

### Supplementary Figures:

a.

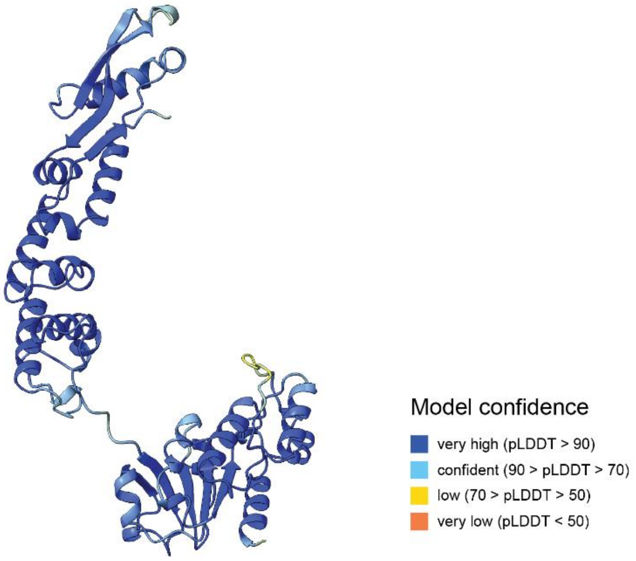

b.

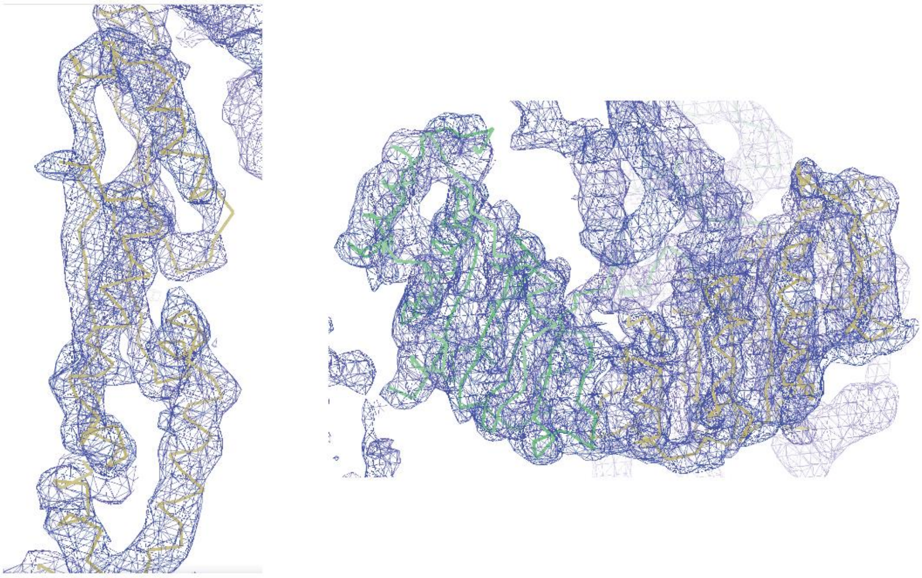

**Supplementary Figure 1. Structural analysis of MksG.** (A) AF model of MksG. The confidence scores of the model are shown on the right. (B) Representative regions of the final electron density map for the three crystal structures contoured at 1.35 $\sigma$, drawn with Coot.

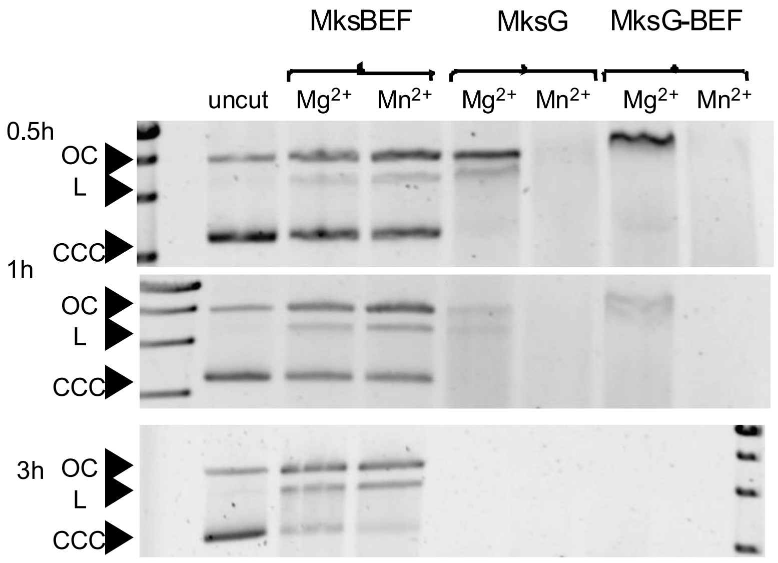

**Supplementary Figure S2: Nicking Assay MksG and MksBEFG.** Nuclease activity assay of MksBEF, MksG and MksBEFG, 10µM (of each) protein were incubated for 0.5, 1, and 3h at 30°C with 250 ng plasmid DNA (pBHK18, 3,337 bp) and 10 mM $Mg^{2+}$ or $Mn^{2+}$. Reactions were stopped by adding 6x purple loading dye (NEB) and boiling samples for 5 min at 90°C. DNA was separated on an agarose gel in TAE buffer

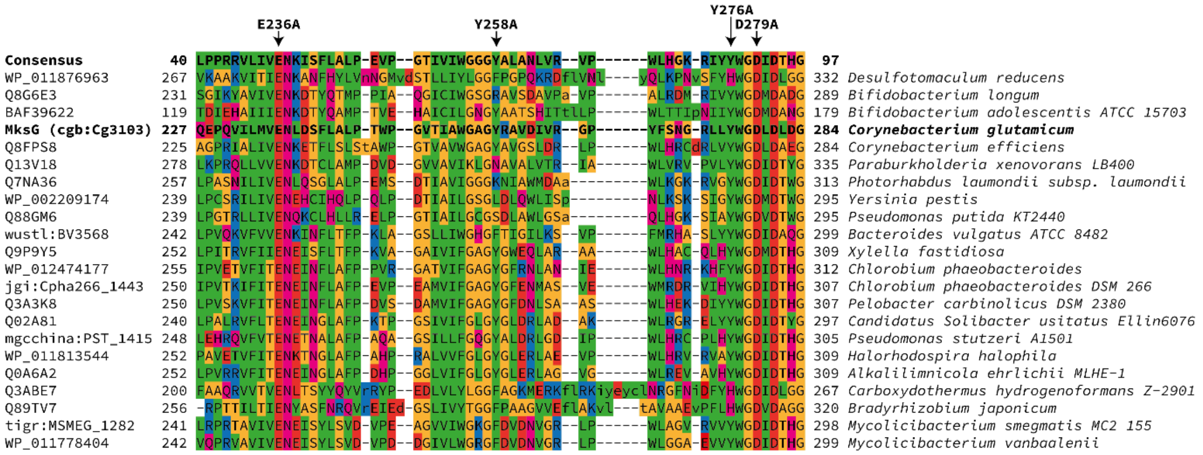

**Supplementary Figure S3: DUF2220 sequence alignment.** Partial visualization of DUF2220 (pfam:09983) sequence conservation alignment (9). The region shown is predicted to be responsible for $Mn^{2+}$ and nucleic acid binding in *C. glutamicum* MksG. Amino acids mutated in the course of this study are indicated via arrows. Amino acids coloring follows the Lesk color scheme.

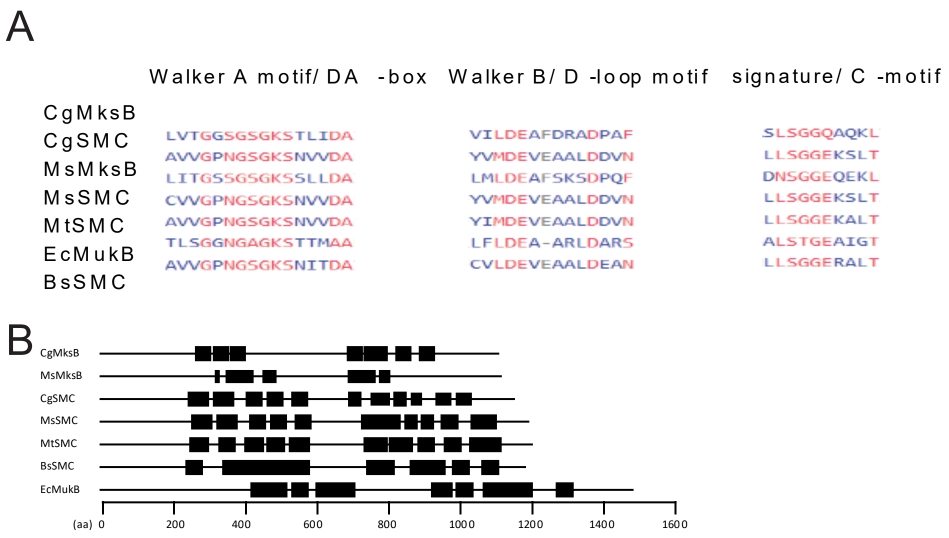

**Supplementary Figure S4: Alignments of sequence motifs in condensin proteins.** (A) Sequence alignment of MksB/SMC/MukB proteins form different organisms, showing the major conserved motifs. Cg, *C. glutamicum*; Ms, *M. smegmatis*; Mt, *M. tuberculosis*; Ec, *E. coli*; Bs, *B. subtlis*. Using NCBI BLASTp (10) (B) Coilel-coild prediction of MksB/SMC/MukB proteins using Coils program (https://mybiosoftware.com/coils-2-2-prediction-coiled-coil-regions-proteins.html) (11). Black squares represent coils. The bottom axis shows the position of the amino acid (aa) corresponding to the coiled-coil segment within a protein's primary structure.

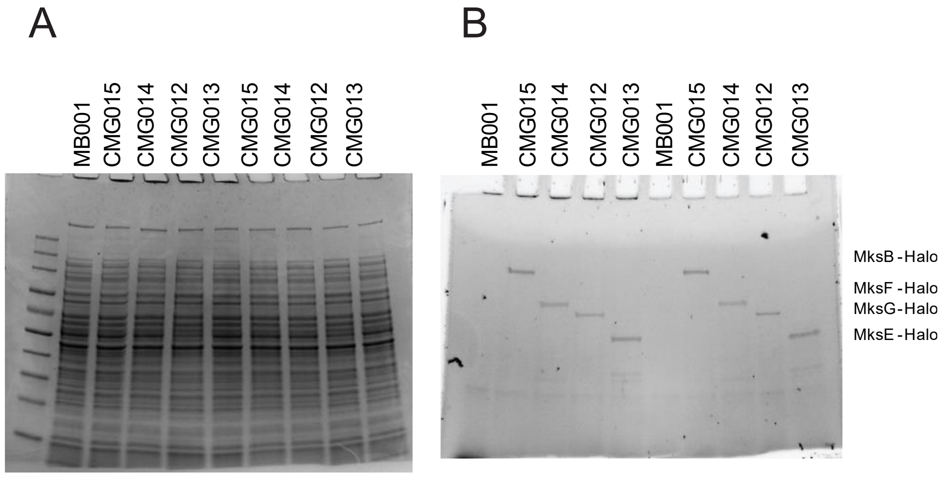

**Supplementary Figure S5: SDS-PAGE analysis of allelic replacement strains with Halo-Tag**. Strains were grown for 5h in BHI medium, $OD_{600}$ was adjusted to OD=15/ml, washed once in PBS buffer. Cells were stained 15 mins with 5 µM TMR dye, washed again once with PBS buffer. Cells were disrupted by sonication. Samples were mixed with 4x SDS-loading dye and heated for 10 min at 40 °C. 20µl/ lane were applied. (A) Coomassie-stained gel; lane 1, marker (NEB, Color pre-stained protein standard), lane 2-6, lysed cells; lane 7-10, cleared lysates. (B) In-gel fluorescence; lane 1-5, lysed cells; lane 6-10, cleared lysates.

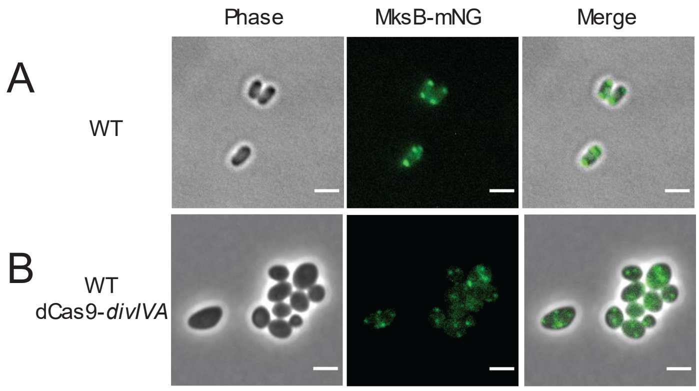

**Supplementary Figure S5: DivIVA-depletion leads to mislocalization of MksB.** Microscopic images of strain (CPF009) expressing MksB-mNeonGreen (mNG) uninduced (A) and CRISPRi/dCas9 for DivIVA-depletion upon induction (B). MksB-mNG foci are shown in green. Scale bar, 2 µm.
